# Identification of 4-(6-((2-methoxyphenyl)amino)pyrazin-2-yl)benzoic acids as CSNK2A inhibitors with antiviral activity and improved selectivity over PIM3

**DOI:** 10.1101/2023.12.04.569845

**Authors:** Kareem A. Galal, Andreas Krämer, Benjamin G. Strickland, Jeffery L. Smith, Rebekah J. Dickmander, Nathaniel J. Moorman, Timothy M. Willson

## Abstract

We report the synthesis of 2,6-disubstituted pyrazines as potent cell active CSNK2A inhibitors. 4’-Carboxyphenyl was found to be the optimal 2-pyrazine substituent for CSNK2A activity, with little tolerance for additional modification. At the 6-position, modifications of the 6-isopropylaminoindazole substituent were explored to improve selectivity over PIM3 while maintaining potent CSNK2A inhibition. The 6-isopropoxyindole analogue **6c** was identified as a nanomolar CSNK2A inhibitor with 30-fold selectivity over PIM3 in cells. Replacement of the 6-isopropoxyindole by isosteric ortho-methoxy anilines, such as **7c**, generated analogues with selectivity for CSNK2A over PIM3 and improved the kinome-wide selectivity. The optimized 2,6-disubstituted pyrazines showed inhibition of viral replication consistent with their CSNK2A activity.

**Graphical Abstract:** 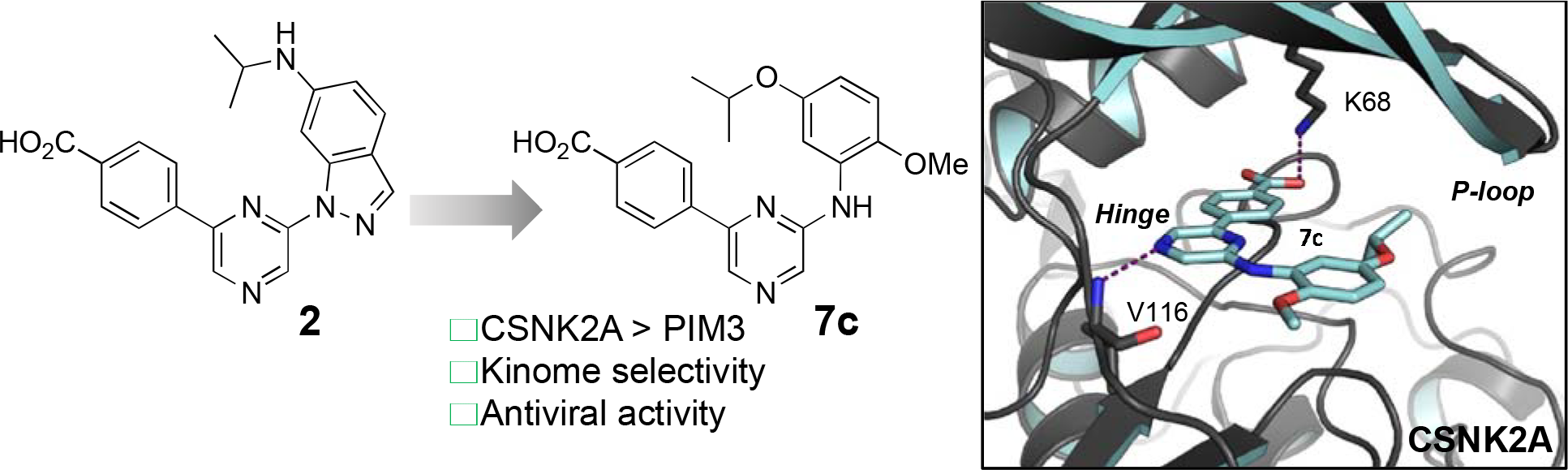

---

Casein Kinase 2 (CSNK2) is member of the eukaryotic protein kinase family that has garnered substantial attention for its multifaceted regulatory role in cell signaling.^1^ CSNK2 is constitutively active and can phosphorylate a wide range of substrates, including transcription factors, cell cycle regulators, and DNA repair proteins. Its dysregulation has been implicated in various pathologies, including cancer, neurological disorders, and autoimmune conditions.^2^ Elevated CSNK2 activity has been observed in many types of cancer, and it is thought to play a role in promoting cell survival, proliferation, and resistance to apoptosis. CSNK2 been explored as a potential anticancer drug target for many years, leading to the clinical development of silmitasertib for the treatment of rare colangiosarcomas.^3^ A distinctive feature of CSNK2 is its tetrameric structure, composed of two catalytic subunits (CSNK2A1 or CSNK2A2), which are the target of ATP-competitive inhibitors, and two regulatory subunits (CSNK2B). Renewed interest in CSNK2 as a potential host target for antiviral therapy followed the discovery of its role in SARS-CoV-2 viral entry.^4^ However, one of the challenges of antiviral drug therapy is the requirement for sustained drug levels at several times the effective antiviral dose. Many of the existing CSNK2A inhibitors (Figure 1) either lack sufficient potency (e.g. silmitisertib)^5^ or good pharmacokinetics (e.g. SGC-CK2-1)^6^ for use as antiviral drugs. Our attempts to modify the naphthyridine chemotype found in silmitisertib failed to yield analogues with increased potency on CSNK2A.^7^ Likewise, extensive modification of the pyrazolopyrimidine chemotype found in SGC-CK2-1 was unable to overcome the high metabolic clearance due to phase 1 and phase 2 metabolism.^8^

**Figure 1.**
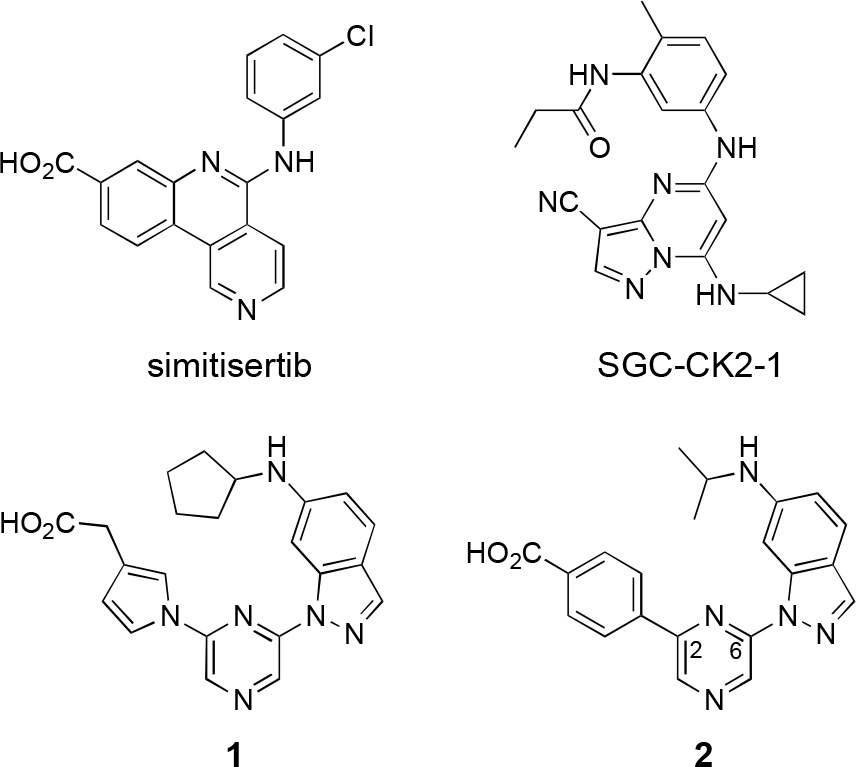
Structures of CSNK2A inhibitors

In the search for an alternative chemotype of CSNK2A inhibitors that would meet the dual requirements of high potency and good pharmacokinetics, we were drawn to the reports by Fuchi^9^ and by Gingipalli^10^ of the pyrazine chemotype (**1** and **2**, Figure 1). Both **1** and **2** are 2,6-disubstituted pyrazines that contain carboxylic acids on their 2-aryl group and a 6-alkylaminoindazole at the 6-position. Pyrazine **1** was reported to be a 9 nM inhibitor of CSNK2A with *in vivo* activity in a rat model of nephritis following interperitoneal dosing.^9^ Pyrazine **2** was described a potent dual CSNK2A/PIM3 inhibitor with IC_50_ of 5 nM and <3 nM, respectively.^10^ Additional characterization of **2** showed that it blocked CSNK2 substrate phosphorylation in cells and showed promising pharmacokinetic properties with low intrinsic clearance, long plasma half-life, and modest oral bioavailability in mice. However, the potent PIM3 kinase inhibition by **2** limited its utility as a pharmacological probe for CSNK2.^10^

The PIM kinases are serine/threonine kinases that regulate cell proliferation, survival, and protein synthesis. There are three highly homologous PIM kinases (PIM1–3), which have similar functions but differential expression across tissues.^11, 12^ Although the ATP-binding sites of the PIM kinases are highly similar, pyrazine **2** had >40-fold selectivity for PIM3 over PIM1 and >460-fold over PIM2.^10^ PIM3 is highly expressed in the lung and its knockdown induces apoptosis of lung cell lines.^13^ Accordingly, the potent PIM3 kinase inhibition of pyrazine **2** would complicate its use as a pharmacological probe to study the role of CSNK2A in lung cells *in vitro* and might lead to toxicity as an antiviral drug *in vivo*. Nevertheless, given its promising pharmacokinetic properties, we elected to synthesize new analogues of the pyrazine chemotype that were optimized in cells for inhibition of CSNK2A and with improved selectivity over PIM3.

A re-synthesized sample of **2** showed an IC_50_ by NanoBRET assay of 12 nM and 18 nM for in-cell target engagement of CSNK2A and PIM3, respectively (Table 1), consistent with the reported values for in vitro enzyme inhibition. Although Gingipalli et al. described the selectivity of analogues of **2** over PIM1 and PIM2, no structure-activity relationship was reported for PIM3.^10^ Therefore, we decided to synthesize analogues with modification of both the 2- and the 6-pyrazine substituents (Figure 1) to establish the determinants of CSNK2A/PIM3 activity and selectivity in cells.

**Table 1.**
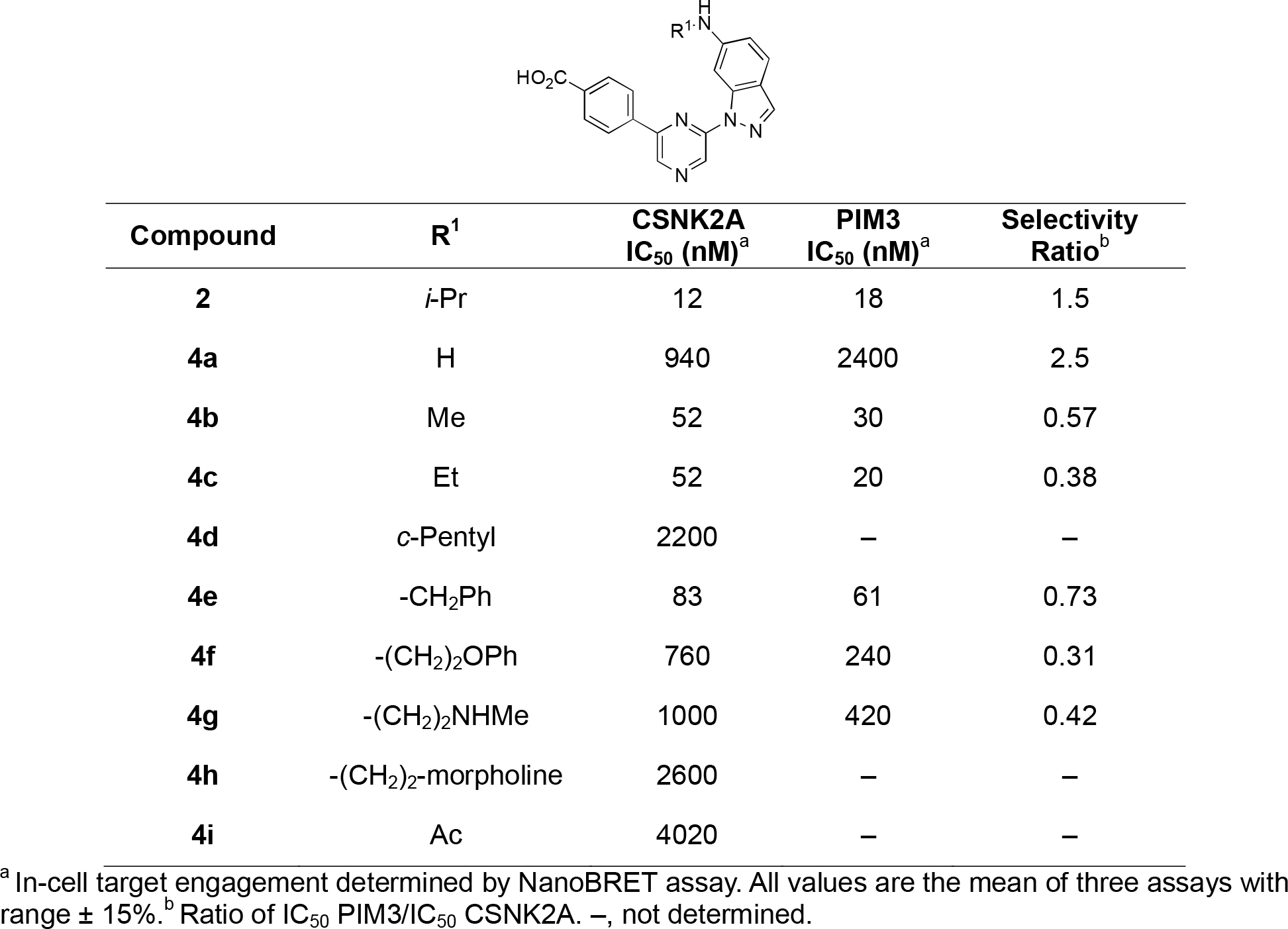
Kinase in-cell target engagement of analogues **2, 4a–i**.

The first series of analogues sought to explore the contribution of the indazole 6-amino substituent. Comparison of the X-ray structure of **1** in CSNK2A with **2** in PIM1 (Figure 2) suggested that subtle differences in conformation of the flexible P-loop might be targeted by this substituent to improve selectivity between the kinases. Synthesis of the modified indazoles started with an S_N_Ar reaction of 6-nitroindazole with 2,6-dichloropyrazine in DMF to produce intermediate **I** in 36% yield (Scheme 1). Reduction of the nitro group using iron and NH_4_Cl afforded intermediate **II** in 88% yield. Palladium catalyzed Suzuki reaction with 4- (carboxymethyl)phenyl boronic acid gave the key intermediate **III** in 79% yield. Saponification of the methyl ester gave the unsubstituted 6-aminoindazole (**4a**). Analogues **4b–e** and **4g** were synthesized by reductive amination of **III** with an aldehyde and NaBH_3_CN, to give esters **3b**-**e**, followed by saponification. Analogues **4f** and **4h** were synthesized by S_N_2 reaction of **III** with an alkyl halide at 80 °C or in a microwave at 170 °C, followed by saponification of the ester. The acetylated amine **4i** was obtained by reaction of **III** with acetyl chloride and subsequent saponification.

**Figure 2.**
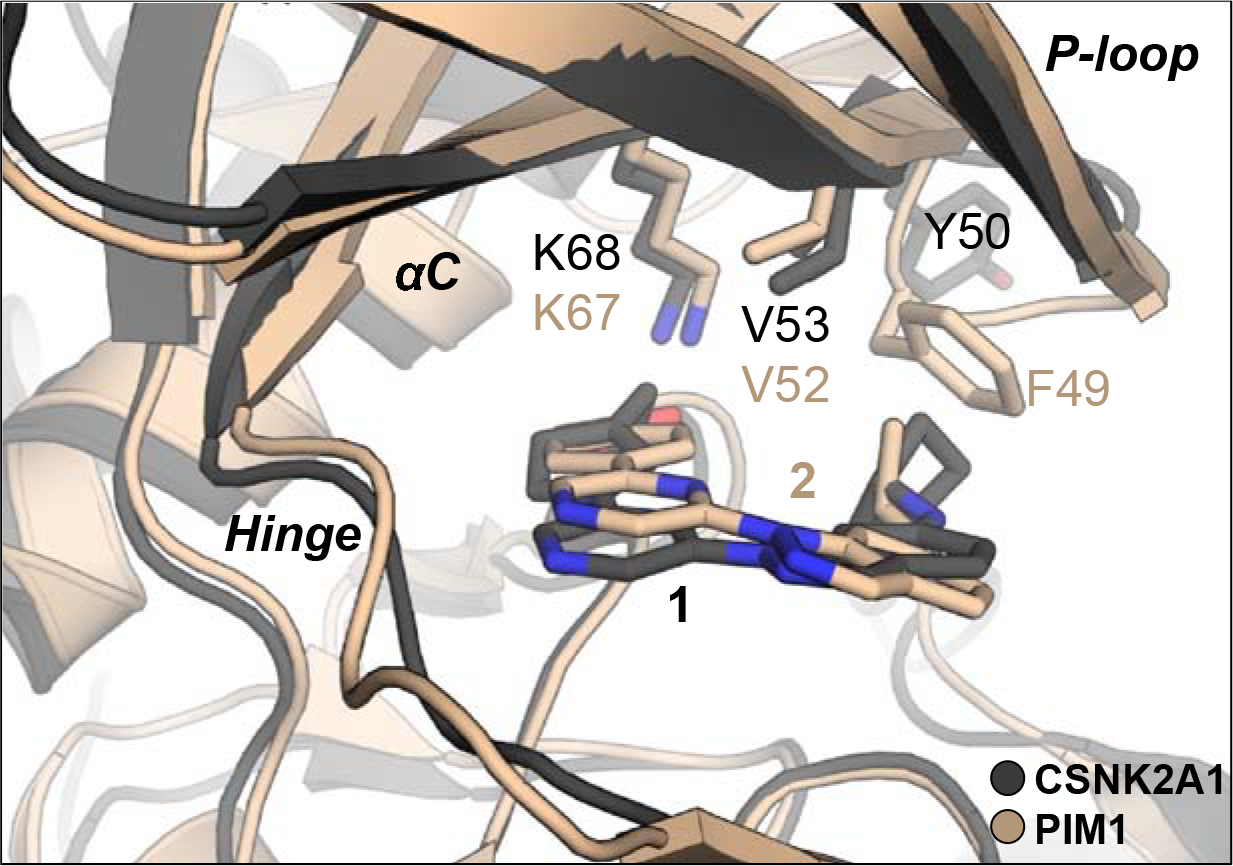
Overlay of X-ray cocrystal structures of **1** in CSNK2A1 (PDB: 3AT3) and **2** in PIM1 (PDB: 6BSK). CSNK2A1 protein with pyrazine **1** are shown in dark grey. PIM1 protein with pyrazine **2** are shown in tan. The carboxylic acids of **1** and **2** form a salt bridge with the catalytic lysine (K68/K67) in both structures. In the flexible P-loop region, V53/V52 interacts with **1** and **2** in both structures but F49 of PIM1 interacts with the isopropylamine of **2** while Y50 of CSNK2A1 is orientated away from the binding pocket with the larger cyclopentylamine of **1**.

**Scheme 1.**
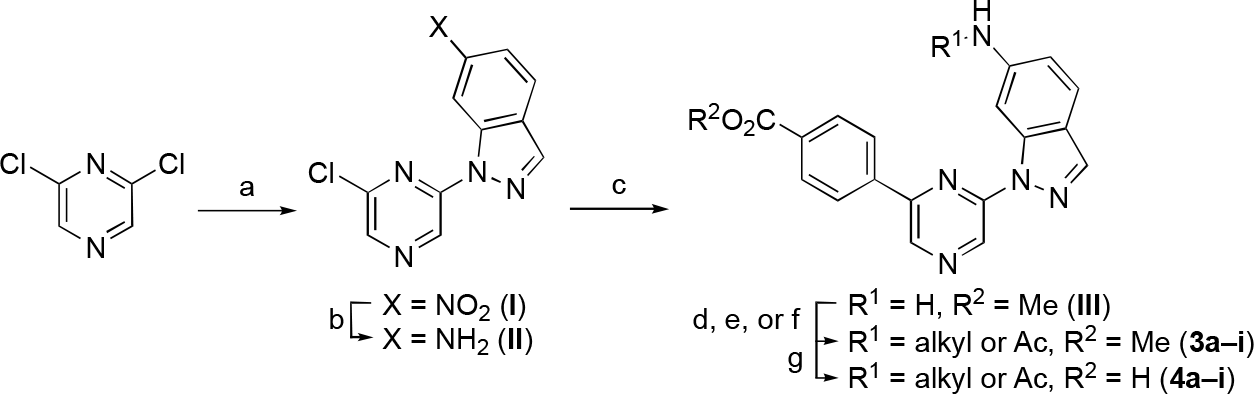
Reagents and conditions: (a) 6-nitro-1*H*-indazole, NaH, DMF, 0°C to rt, 32%; (b) Fe, NH_4_Cl, EtOH/water, reflux, 88%; (c) 4-(methoxycarbonyl) phenyl boronic acid, Pd(dppf)Cl_2_,Na_2_CO_3_ toluene/EtOH/water, 70 °C, 79%; (d) aldehyde, NaBH_3_CN, THF, 41-43%; (e) alkyl halide, KI, K_2_CO_3_, ACN, 170 °C, MW (<u>or</u> 80 °C with no MW), 12-40%; (f) acetyl chloride, DIPEA, THF, 0 °C to rt, 41%; (g) 2 M LiOH, THF, rt, 9–98%.

The analogues **4a–i** were evaluated for their kinase in-cell target engagement using NanoBRET assays^14^ developed for CSNK2A and PIM3 (Table 1). For the CSNK2A NanoBRET we opted to use the CSNK2A2 isoform since it gave a larger assay window and the ATP-binding site is identicial to CSNK2A1. The isoproylamine **2** was a potent inhibitor of both kinases with a slight preference for CSNK2A over PIM3. Removal isopropyl group in **4a** resulted in a large decrease in activity on both kinases. Addition of a methyl (**4b**) or ethyl (**4c**) group restored kinase inhibition, but both analogues were more potent on PIM3 than CSNK2A suggesting that the branched alkyl group of **2** was preferable. Unfortunately, the larger cyclo-pentyl group (**4d**) was not tolerated by CSNK2A in this series. The result was surprising given the prior report of potent CSNK2A enzyme inhibition of **1** which also contains a 6-cyclopentylaminoindazole.^9^ The benzyl analogue (**4e**) showed good activity on both kinases but still showed a preference for PIM3. The larger phenoxyethyl (**4f**) analogue was less active. Addition of a second basic group in **4g** and **4h** gave analogues with poor activity on CSNK2A. The acylamine (**4i**) had only weak activity on CSNK2A.

Since these initial results suggested that the 6-isopropylaminoindazole was the optimal 6-pyrazine substituent for CSNK2A activity, we designed a second series of analogues with modification of the 2-pyrazine benzoic acid group to explore the effect on CSNK2/PIM3 selectivity. Modification of the benzoic acid required a different synthesis (Scheme 2) which started from commercially available 6-aminoindazole. Reductive amination with acetone and NaBH(OAc)_3_ gave intermediate **IV** containing the isopropylamine in 70–88% yield. Ulmann-type coupling with 2,6-diiodopyrazine using (1*R*, 2*R*)-*N,N*-dimethylcyclohexane-1,2-diamine as a ligand and CuI as a catalyst at 110 °C generated intermediate **V** in 30–45% yield. Analogues **5a–e** were synthesized by Suzuki coupling with an arylboronic acid and Pd(dppf)Cl_2_ in 45– 67%yield, followed by saponification of the ester if required. To synthesize the 3-chloroindazole analogue (**5f**), the sequence was repeated from 3-choloro-6-aminoindazole as a starting material (Scheme 2).

Analogues **5a–f** were tested for kinase in-cell target engagement by NanoBRET assay (Table 2). Replacement of the benzoic acid with the corresponding carboxamide (**5a**) or sulfonamide (**5b**) resulted in analogues that were inactive on CSNK2A. In the analogues that retained the benzoic acid, addition of a 3-fluoro group (**5c**) restored CSNK2A activity but resulted in a compound that had no selectivity over PIM3. The larger 3-chloro group in **5d** resulted in a decrease in CSNK2A activity and the 3-methoxy analogue (**5e**) was inactive. Analogue **5f** with a 3-chloro group on the indazole core had potent activity on both CSNK2A and PIM3 but showed no selectivity between the kinases. However, **5f** demonstrated that modification of the indazole at the pyrazine 6-position was better tolerated than changes to the benzoic acid substituent.

**Table 2.**
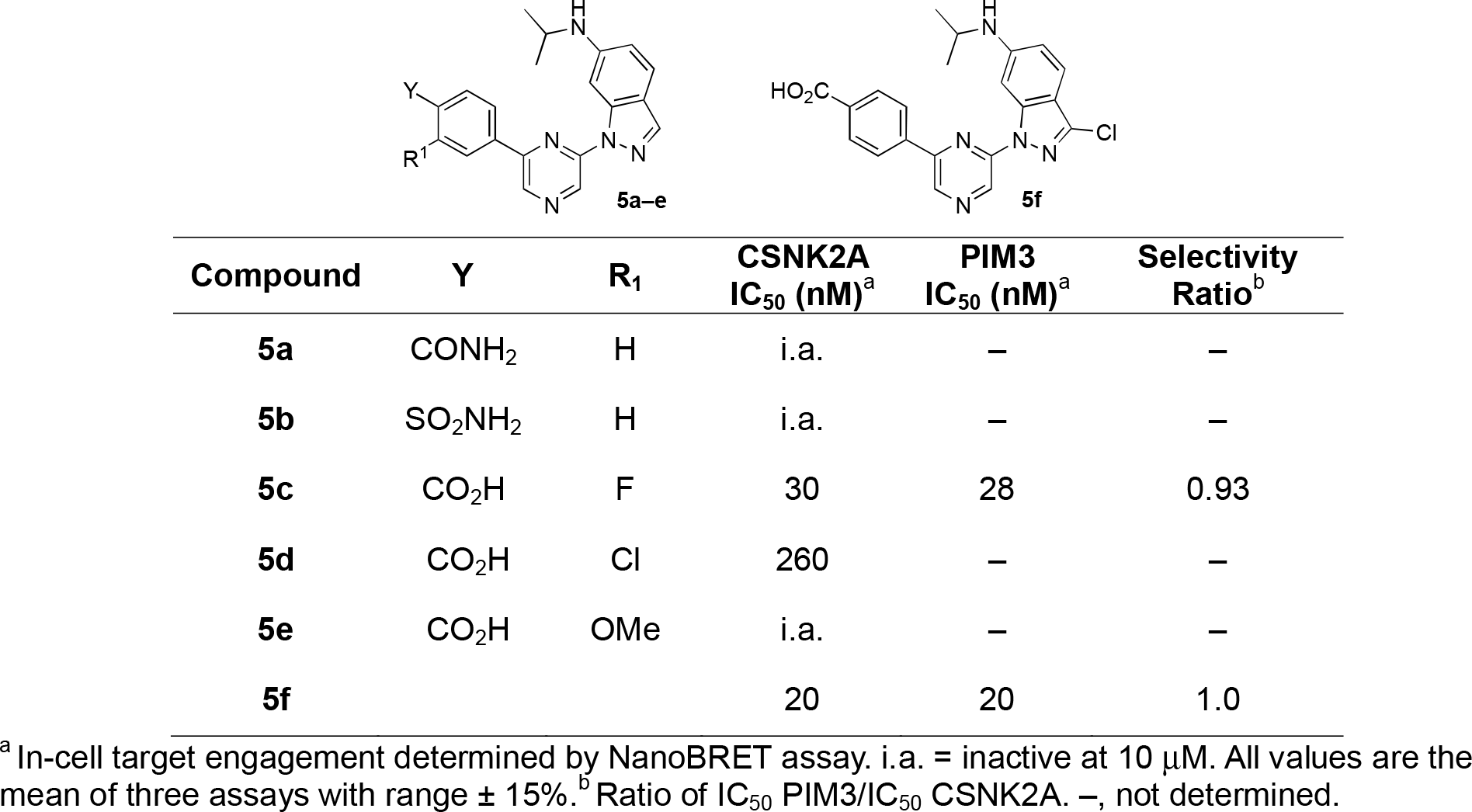
Kinase in-cell target engagement of analogues **5a–f**.

**Scheme 2.**
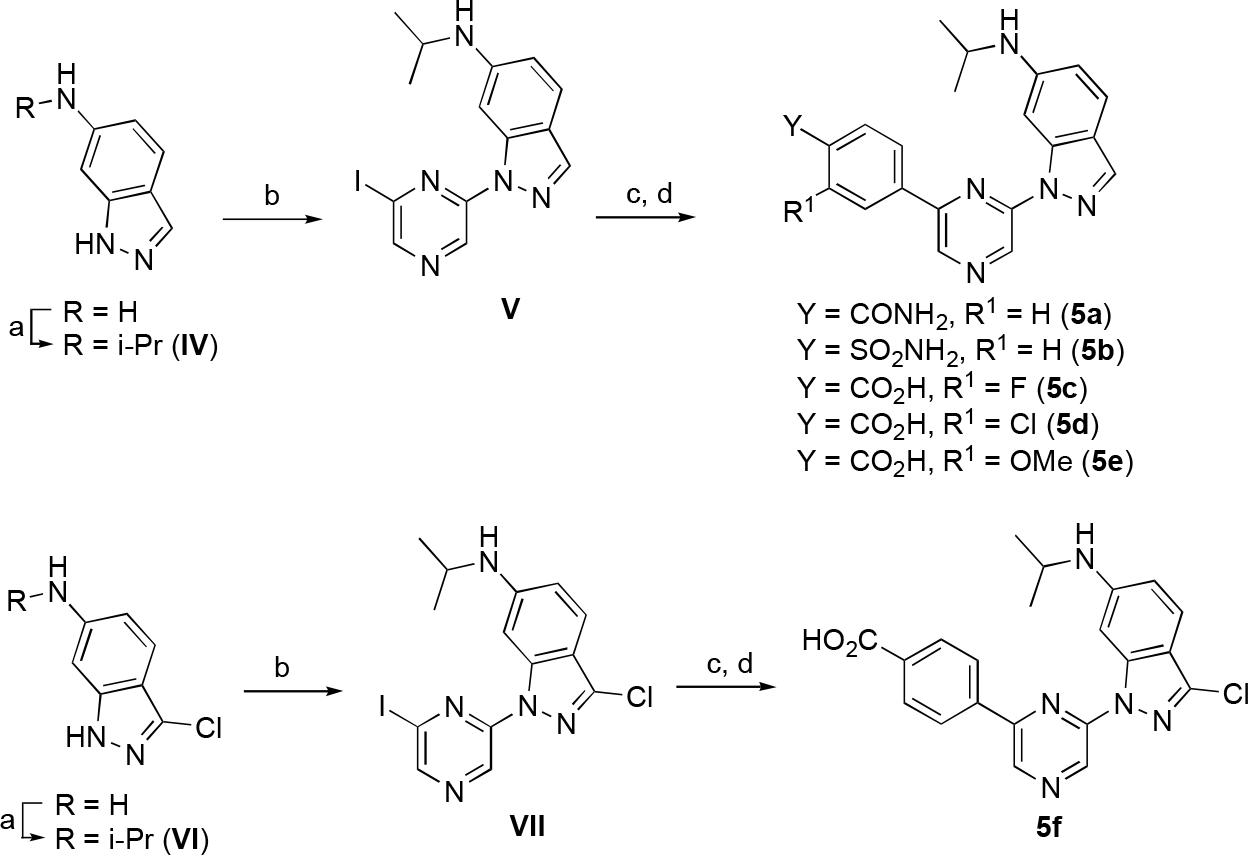
Reagents and conditions: (a) acetone, AcOH, NaBH(OAc)_3_, DCM, rt, 70–88%; (b) 2,6-diiodopyrazine (1*R*, 2*R*)-*N*1,*N*2-dimethylcyclohexane-1,2-diamine, CuI, K_3_PO_4_, dioxane, 110 °C, 24 h, 30–45%; (c) boronic acid, Pd(dppf)Cl_2_, Na_2_CO_3_, toluene/EtOH/water, 70 °C, 45–67%; (d) 2 M LiOH, THF, rt, 22–62%

A third series of analogues was synthesized in which the indazole 6-amino substituent was replaced by an ether. To synthesize the ether analogue of **2**, 6-hydroxyindazole was alkylated with 2-iodopropane in 63% yield to give intermediate **VIII** (Scheme 3). A Ulman coupling reaction with 2,6-diiodopyrazine using (1*R*, 2*R*)-*N,N*-dimethylcyclohexane-1,2-diamine and CuI at 110 °C gave a modest 31% yield of **IX**. Suzuki reaction with (4-(methoxycarbonyl) phenyl)boronic acid followed by ester hydrolysis produced isopropyl ether **6a**. Synthesis of the methoxy ether analogue (**6b**) started with an S_N_Ar reaction between 2,6-dichloropyrazine and 6-methoxyindazole to give intermediate **X** in 50% yield. A repeat of the Suzuki coupling and ester hydrolysis yielded **6b**. The indole analogue of **6a** was synthesized by reaction of 2-chloro-6-bromopyrazine with (4-(methoxycarbonyl)phenyl)boronic acid by Suzuki coupling using Pd(PPh_3_)_4_ as catalyst to give intermediate **XI** in 49% yield. An S_N_Ar reaction with 6-isoproxyindole using sodium hydride as a base produced indole **6c** in low yield. No saponification was required for this analogue since the ester was removed during the S_N_Ar reaction. The 6-isopropylaminoindole **6d** was synthesized from the corresponding indole by S_N_Ar reaction, followed by saponification.

**Scheme 3.**
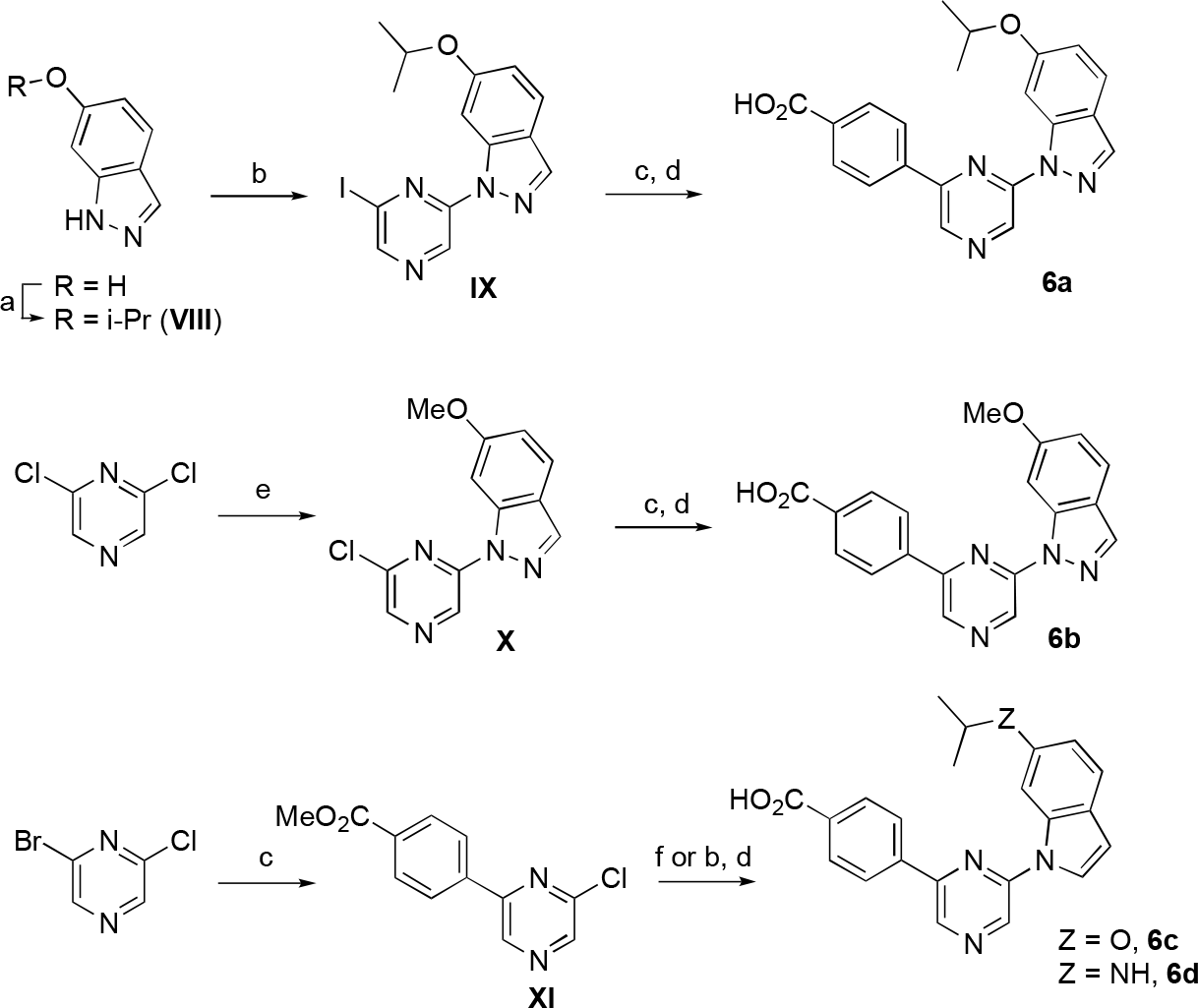
Reagents and conditions: (a) 2-iodopropane, Cs_2_CO_3_, DMF, rt, 63%; (b) 2,6-diiodopyrazine (1*R*, 2*R*)-*N,N*-dimethylcyclohexane-1,2-diamine, CuI, K_3_PO_4_, dioxane, 110 °C, 24 h, 13–31%; (c) 4- (methoxycarbonyl)phenylboronic acid, Pd(dppf)Cl_2_ (or Pd(PPh_3_)_4_, Na_2_CO_3_, toluene/EtOH/water, 70 °C, 32–61%; (d) 2 M LiOH, THF, rt, 52–65%; (e) 6-methoxyindazole, NaH, DMF, 0 °C to rt, 50%; (f) 6-isoproxyindole or 6-isopropylaminoindole, NaH, DMF, 0 °C to rt, 10%.

Analogues **6a–c** demonstrated nanomolar activity for in-cell target engagement on CSNK2A and PIM3 by NanoBRET assay (Table 3). The isopropoxyether **6a** had equivalent activity to **2** in cells, demonstrating that the 6-anilinoindazole nitrogen was not required for potent inhibition of either CSNK2A or PIM3. The methyl ether **6b** was less active on CSNK2A but maintained its activity on PIM3 suggesting that there were indeed subtle differences in the P-loop of their ATP binding pockets. Most surprisingly, switching the 6-isopropoxyindazole to the 6-isopropoxyindole **6c** resulted in an increase in CSNK2A activity and a concomitant decrease in PIM3 activity. 6-isopropoxyindole **6c** was the most potent and selective analogue in the pyrazine series, demonstrating a 30-fold preference for CSNK2A over PIM3. The 6-isopropylaminoindole (**6d**) also showed potent CSNK2 inhibition with reduced PIM3 activity with a 13-fold selectivity ratio.

**Table 3.** Kinase in-cell target engagement of analogues **6a–c**.

| Compound | R<sup>1</sup> | CSNK2A<br>IC<sub>50</sub> (nM)<sup>a</sup> | PIM3<br>IC<sub>50</sub> (nM)<sup>a</sup> | Selectivity<br>Ratio<sup>b</sup> |
| --- | --- | --- | --- | --- |
| **6a** | O*i*-Pr | 11 | 11 | 1 |
| **6b** | OMe | 97 | 11 | 0.11 |
| **6c** | O*i*-Pr | 2.2 | 66 | 30 |
| **6d** | N*i*-Pr | 18 | 230 | 13 |

<sup>a</sup> In-cell target engagement determined by NanoBRET assay. All values are the mean of three assays with range $\pm 15\%$ . <sup>b</sup> Ratio of IC<sub>50</sub> PIM3/IC<sub>50</sub> CSNK2A

Since the switch to the indole in **6c** and **6d** had produced the most selective analogues, we decided to synthesize several isosteric replacements of the heterocycle. Suzuki *et al*. had reported that meta-substituted aniline replacements of the indazole in **1** retained CSNK2A inhibition,^15^ but did not synthesize any 2-substituted anilines that might retain the planarity of an indole through H-bonding. We designed anilines with either fluorine or methoxy as an ortho-substituent to provide a potential H-bond acceptor for the aniline N-H (Scheme 4). To synthe-size the compounds, 2-fluoro-5-nitroaniline or 2-methoxy-5-nitroaniline was reacted with intermediate **XI** by Buchwald amination reaction, to afford intermediate **XIIa** and **XIIb** in 45–55% yield. The resulting nitro intermediates were reduced with Fe and NH_4_Cl to give the corresponding anilines **XIIIa** and **XIIIb**. Reductive amination with acetone and saponification yielded the isopropyl anilines **7a** and **7b**. Synthesis of the isopropyl ether **7c** started from 3-nitro-4-methoxyphenol. Reaction with 2-iodopropane produced the ether **XIV** in 70% yield. Reduction of the nitro group to the intermediate **XVI** followed by Buchwald coupling with intermediate **XI** and saponification yielded the isopropyl ether analogue **7c**.

**Scheme 4.**
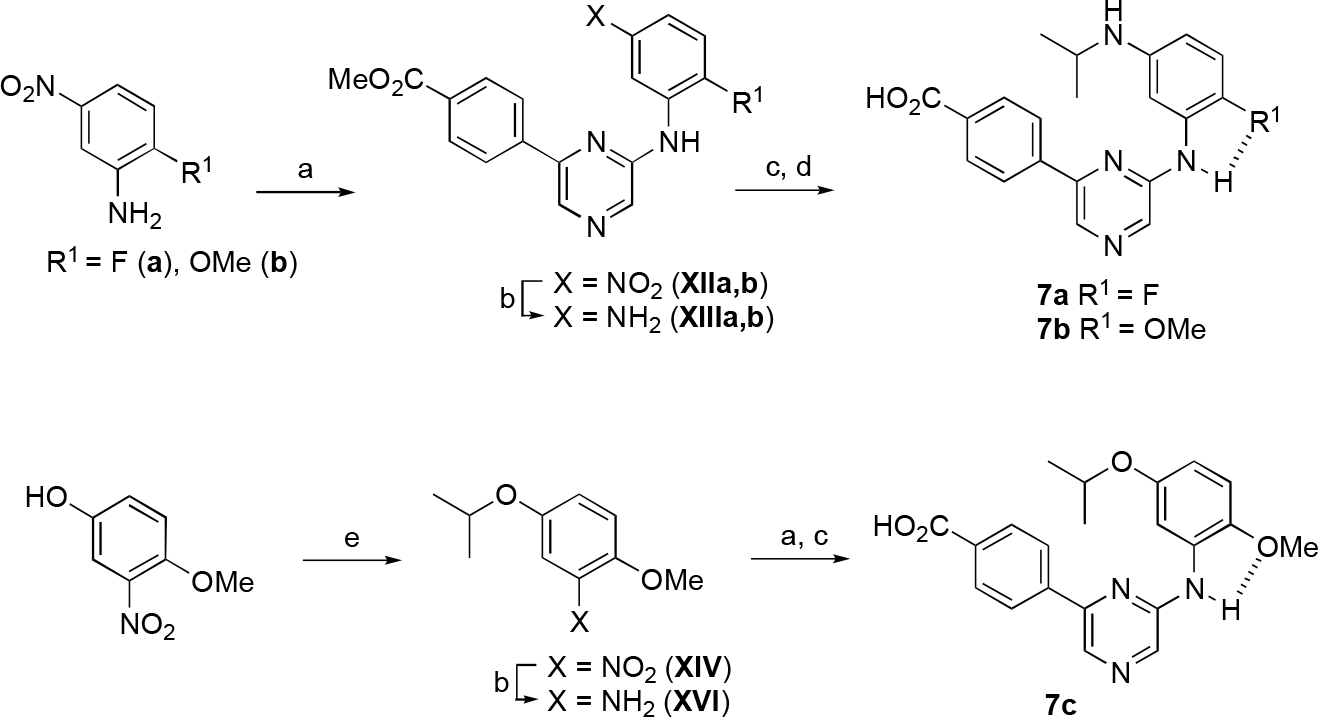
Reagents and conditions: **XI**, Pd_2_(dba)_3_, BINAP, Cs_2_CO_3_, dioxane, 90 °C, 45-55%; (b) Fe, NH_4_Cl, EtOH/water, reflux; (c) acetone, NaBH_3_CN, AcOH, DMF, rt, 26-33%; (d) 2 M LiOH, THF, rt, 19-93%; (e) 2-iodopropane, Cs_2_CO_3_, DMF, rt, 70%. Dashed lines show the potential intramolecular H-bonds.

The three aniline analogues **7a–c** were assayed by NanoBRET for in-cell target engagement to CSNK2A and PIM3 (Table 4). The 2-fluoroaniline **7a** maintained moderate activity on CSNK2A but showed a reduction in PIM3 activity. As a result, **7a** was weakly selective for CSNK2A over PIM3. The 2-methoxyaniline **7b** had equivalent activity to **7a** on CSNK2A but with an even greater reduction in PIM3 activity. The selectivity of aniline **7b**, with a ratio of 20-fold, rivaled that of indole **6c**. Switching the isopropyl aniline to an isopropyl ether in **7c** resulted in an increase in both CSNK2A and PIM3 activity, with the compound showing modest selectivity for CSNK2A (5.6-fold).

**Table 4.** Kinase in-cell target engagement of analogues **7a–c**.

| Compound | R<sup>1</sup> | Z | CSNK2A<br>IC<sub>50</sub> (nM)<sup>a</sup> | PIM3<br>IC<sub>50</sub> (nM)<sup>a</sup> | Selectivity<br>Ratio<sup>b</sup> |
| --- | --- | --- | --- | --- | --- |
| **7a** | F | N | 120 | 430 | 3.6 |
| **7b** | OMe | N | 100 | 2000 | 20 |
| **7c** | OMe | O | 60 | 340 | 5.6 |

<sup>a</sup> In-cell target engagement determined by NanoBRET assay. All values are the mean of three assays with range ± 15%. <sup>b</sup> Ratio of IC<sub>50</sub> PIM3/IC<sub>50</sub> CSNK2A

The indazole **2**, indole **6c** and ortho-methoxy aniline **7c** were also evaluated across a large panel of 101 kinases by thermal shift assays to identify other potential off-target kinases (Table S1). Aniline **7c** showed activity on 18/101 kinases with T_m_ >5 °C at a concentration of 10 µM. The major off target kinases with T_m_ >9 °C were DAPK3, PIM1, BIKE, MAPK15, and DYRK2. Notably, the indazole **2** and indole **6c** were significantly less selective across the panel with 34 and 29 kinases showing T_m_ >5 °C, respectively. Thus, the ortho-methoxy aniline functioned as a isosteric replacement for the indazole and indole that maintained submicromolar activity on CSNK2A in cells and also improved the broader kinase selectivity.

To provide additional insight onto the basis of the potent kinase inhibition of the 6-indolo-pyrazine **6c** and the 6-anilino-pyrazine **7c**, we determined their co-crystal structures with CSNK2A (Figure 3). The structure of indole **6c** (PDB: 8QWY) showed the expected interaction within the ATP-binding pocket of CSNK2A. The carboxylic acid interacted with the catalytic lysine (K68), while one of the pyrazine nitrogens formed the main H-bond with the hinge region (Figure 3A). The three rings of **6c** adopted a co-planar conformation as is commonly seen in other CSNK2A inhibitors. The aniline **7c** (PDB: 8QWZ) adopted a similar conformation in the CSNK2A ATP-binding pocket, with polar interactions to K68 and the hinge region. The three rings of **6c** were also co-planar, supported in part by an internal H-bond between the aniline N-H and the ortho-methoxy group (Figure 3B). The co-crystal structures confirmed the effectiveness of the ortho-methoxy aniline **7c** as an isosteric replacement of the indole **6c**.

**Figure 3.**
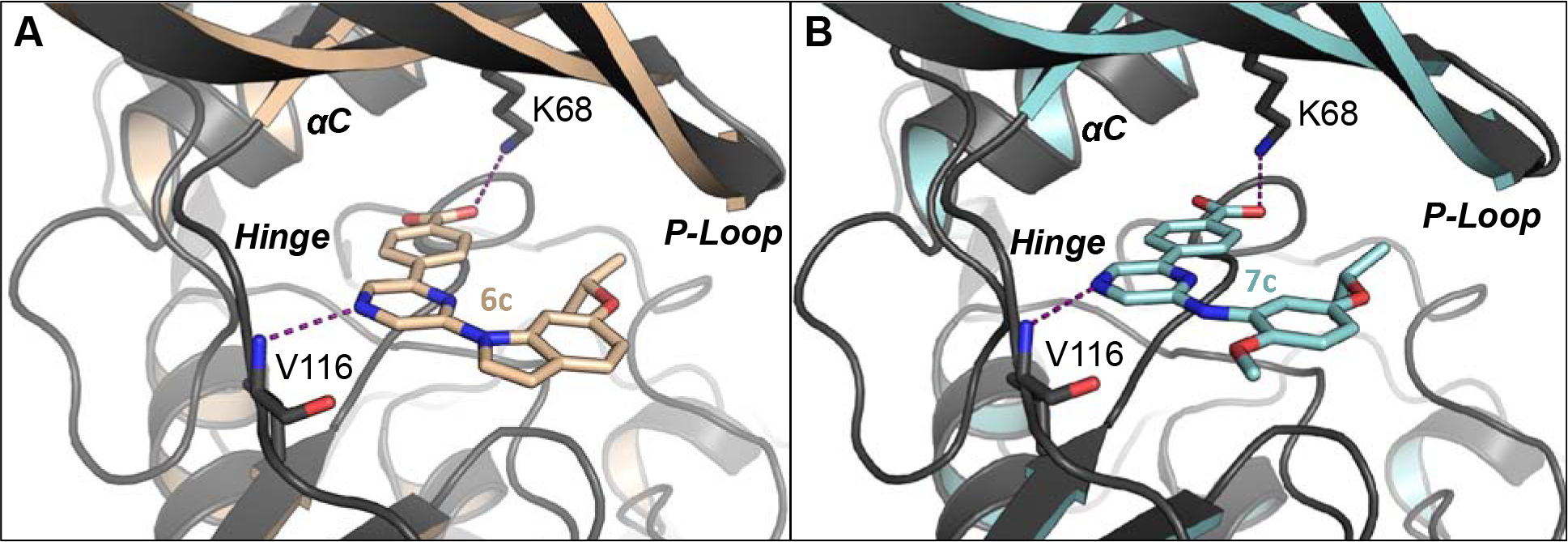
CSNK2A X-ray co-crystal structures. (A) Indole **6c** (PDB: 8QWY); (B) Aniline **7c** (PDB: 8QWZ); Polar interactions between the carboxylic acid and K68 and the pyrazine nitrogen and the backbone amide of V116 are shown as dashed lines.

The pyrazines with IC_50_ <100 nM on CSNK2A and the unsubstituted indazole **4a** were assayed for antiviral activity against the murine hepatitis virus (MHV),^16^ a non-pathogenic β-coronavirus that is orthologous to SARS-CoV-2 (Table 5). The pyrazines displayed IC_50_ ranging from 0.8–8.6 µM for inhibition of viral replication. Although many of the pyrazines showed either no or modest selectivity for CSNK2A over PIM3, a plot of kinase IC_50_ against antiviral activity showed that the potency for MHV inhibition tracked better with CSNK2A activity than PIM3 (Figure 4).

**Table 5.** Antiviral activity of pyrazine CSNK2A inhibitors.

| Compound | MHV IC<sub>50</sub> (μM)<sup>a</sup> |
| --- | --- |
| **2** | 1.1 |
| **4a** | 4.4 |
| **4b** | 8.6 |
| **4c** | 1.2 |
| **4e** | 6.3 |
| **5c** | 3.0 |
| **5f** | 2.2 |
| **6a** | 3.2 |
| **6b** | 1.0 |
| **6c** | 0.8 |
| **6d** | 0.7 |
| **7a** | 2.3 |
| **7b** | 1.8 |
| **7c** | 1.0 |

<sup>a</sup> Inhibition of replication of MHV-nLuc in DBT cells

**Figure 4.**
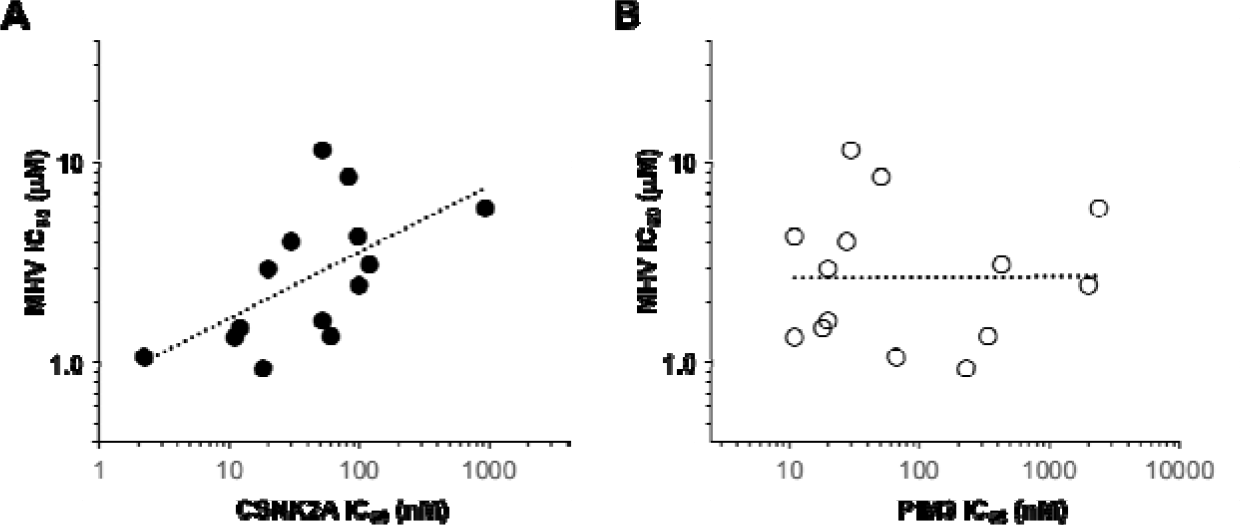
Relationship of kinase inhibition and antiviral activity. (A) CSNK2A in-cell target engagement predicts MHV inhibition with *R* = 0.4; (B) PIM3 in-cell target engagement did not correlate with MHV inhibition.

In conclusion, 2,6-disubstitued pyrazines are a promising chemotype of CSNK2A inhibitors. The optimal 4-carboxyphenyl substituent at the pyrazine 2-position can be readily synthesized by palladium catalyzed Suzuki coupling. At the pyrazine 6-position, incorporation of 6-isopropoxyindole resulted in an analogue **6c** with nanomolar in-cell target engagement for CSNK2A and 30-fold selectivity over PIM3. The ortho-methoxy aniline **7c** also showed potent in-cell target engagement for CSNK2A and had improved kinome-wide selectivity by thermal shift assays. The demonstration that ortho-substituted anilines were effective bioisoteric replacements for the 6-isopropoxyindole highlights the potential for modular synthesis of new 2,6-disubstitued pyrazine CSNK2A inhibitors optimized for cellular potency and antiviral activity.

## Abbreviations

ACN: acetonitrile
ATP: adenosine triphosphate
BRET: bioluminescence resonance energy transfer
DCM: dichloromethane
DIPEA: diisopropylethylamine
DMF: dimethylformamide
DNA: deoxyribonucleic acid
HEK: human embryonic kidney

## Acknowledgments

The Structural Genomics Consortium (SGC) is a registered charity (no: 1097737) that receives funds from Bayer AG, Boehringer Ingelheim, Bristol Myers Squibb, Genentech, Genome Canada through Ontario Genomics Institute [OGI-196], EU/EFPIA/OICR/McGill/KTH/Diamond Innovative Medicines Initiative 2 Joint Undertaking [EUbOPEN grant 875510], Janssen, Merck KGaA (aka EMD in Canada and US), Pfizer and Takeda. Research reported in this publication was supported in part by the NC Biotech Center Institutional Support Grant 2018-IDG-1030, by the NIH Illuminating the Druggable Genome 1U24DK116204-01. This project was supported by the Rapidly Emerging Antiviral Drug Development Initiative (READDI) at the University of North Carolina at Chapel Hill with funding from the North Carolina Coronavirus State and Local Fiscal Recovery Funds program, appropriated by the North Carolina General Assembly.

## Supplementary Material

Supplementary data associated with this article can be found, in the online version.

## References

1. Litchfield DW. Protein kinase CK2: structure, regulation and role in cellular decisions of life and death. Biochem J. 2003;369(Pt 1): 1–15.

2. Borgo C, D’Amore C, Sarno S, Salvi M, Ruzzene M. Protein kinase CK2: a potential therapeutic target for diverse human diseases. Signal Transduct Target Ther. 2021;6(1): 183.

3. Strum SW, Gyenis L, Litchfield DW. CSNK2 in cancer: pathophysiology and translational applications. Br J Cancer. 2022;126(7): 994–1003.

4. CP QM, M. R. Protein Kinase CK2 and SARS-CoV-2: An Expected Interplay Story. Kinases Phosphatases. 2023;1(2): 141–150.

5. Yang X, Dickmander RJ, Bayati A, et al. Host Kinase CSNK2 is a Target for Inhibition of Pathogenic SARS-like β-Coronaviruses. ACS Chem Biol. 2022;17(7): 1937–1950.

6. Wells CI, Drewry DH, Pickett JE, et al. Development of a Potent and Selective Chemical Probe for the Pleiotropic Kinase CK2. Cell Chem Biol. 2021;28(4): 546–558 e510.

7. Davis-Gilbert ZW, Kramer A, Dunford JE, et al. Discovery of a Potent and Selective Naphthyridine-Based Chemical Probe for Casein Kinase 2. ACS Med Chem Lett. 2023;14(4): 432–441.

8. Yang X, Ong HW, Dickmander RJ, et al. Optimization of 3-Cyano-7-cyclopropylamino-pyrazolo[1,5-a]pyrimidines Toward the Development of an In Vivo Chemical Probe for CSNK2. bioRxiv. 2023.

9. Fuchi N, Iura Y, Kaneko H, et al. Discovery and structure-activity relationship of 2,6-disubstituted pyrazines, potent and selective inhibitors of protein kinase CK2. Bioorg Med Chem Lett. 2012;22(13): 4358–4361.

10. Gingipalli L, Block MH, Bao L, et al. Discovery of 2,6-disubstituted pyrazine derivatives as inhibitors of CK2 and PIM kinases. Bioorg Med Chem Lett. 2018;28(8): 1336–1341.

11. Narlik-Grassow M, Blanco-Aparicio C, Carnero A. The PIM family of serine/threonine kinases in cancer. Med Res Rev. 2014;34(1): 136–159.

12. Santio NM, Koskinen PJ. PIM kinases: From survival factors to regulators of cell motility. Int J Biochem Cell Biol. 2017;93: 74–85.

13. Fan X, Xie Y, Zhang L, et al. Effect of Pim-3 Downregulation on Proliferation and Apoptosis in Lung Adenocarcinoma A549 Cells. Ann Clin Lab Sci. 2019;49(6): 770–776.

14. Vasta JD, Corona CR, Wilkinson J, et al. Quantitative, Wide-Spectrum Kinase Profiling in Live Cells for Assessing the Effect of Cellular ATP on Target Engagement. Cell Chem Biol. 2018;25(2): 206–214 e211.

15. Suzuki Y, Cluzeau J, Hara T, et al. Structure-activity relationships of pyrazine-based CK2 inhibitors: synthesis and evaluation of 2,6-disubstituted pyrazines and 4,6-disubstituted pyrimidines. Arch Pharm (Weinheim). 2008;341(9): 554–561.

16. Korner RW, Majjouti M, Alcazar MAA, Mahabir E. Of Mice and Men: The Coronavirus MHV and Mouse Models as a Translational Approach to Understand SARS-CoV-2. Viruses. 2020;12(8): 880.

