## Supporting Information for "Identification of 4-(6-((2-methoxyphenyl)amino)pyrazin-2-yl)benzoic acids as CSNK2A inhibitors with antiviral activity and improved selectivity over PIM3"

**Table of Contents** **Page**

Table S1. Kinase selectivity by thermal shift assay S2
Chemistry Methods S3
Biology Methods S19

**Table S1**. Kinase selectivity of **2**, **6c**, and **7c** by thermal stability assay. Compounds were tested at a concentration of 10 μM. $T_m$ shifts are shown in ºC. Kinases with a $T_m$ shift >5 ºC are highlighted in red.

| Kinase | 2 | 6c | 7c |
|---|---|---|---|
| DAPK3 | 16 | 11 | 14 |
| CSNK2a2 | 15 | 16 | 15 |
| CSNK2a1 | 15 | 15 | 15 |
| PIM1 | 13 | 10 | 9 |
| BIKE | 13 | 11 | 10 |
| MAPK15 | 12 | 15 | 10 |
| DYRK2 | 12 | 11 | 9 |
| GAK | 11 | 9 | 7 |
| MSK1_b | 11 | 6 | 6 |
| PLK4 | 11 | 10 | 6 |
| AurA | 10 | 8 | 4 |
| PIM3 | 9 | 10 | 6 |
| DAPK1 | 9 | 7 | 6 |
| LOK | 8 | 4 | 4 |
| HIPK2 | 8 | 9 | 7 |
| CLK3 | 8 | 8 | 7 |
| PHKg2 | 8 | 5 | 3 |
| DRAK1 | 8 | 6 | 4 |
| ABL1 | 7 | 8 | 7 |
| AAK1 | 7 | 6 | 5 |
| FLT1 | 7 | 7 | 5 |
| ULK3 | 7 | 5 | 3 |
| CLK1 | 7 | 7 | 7 |
| DYRK1A | 7 | 6 | 5 |
| GPRK5 | 6 | 4 | 4 |
| MELK | 6 | 4 | 4 |
| SRPK1 | 6 | 6 | 3 |
| CHK2 | 6 | 4 | 3 |
| JNK1 | 6 | 3 | 1 |
| BRAF | 6 | 8 | 7 |
| BMPR2 | 6 | 7 | 0 |
| JNK3 | 6 | 3 | 1 |
| MAP3K5 | 5 | 3 | 1 |
| MEK4 | 5 | 9 | 2 |

| Kinase | 2 | 6c | 7c |
|---|---|---|---|
| DCAMKL1 | 4 | 3 | 3 |
| MST2 | 4 | 3 | 2 |
| CDK2 | 4 | 5 | 1 |
| AURKB | 4 | 4 | 2 |
| RSK1_b | 4 | 2 | 1 |
| ULK1 | 3 | 2 | 1 |
| MER | 3 | 3 | 3 |
| MST1 | 3 | 1 | 1 |
| SLK | 3 | 2 | 1 |
| CK1d | 3 | 3 | 2 |
| SRC | 3 | 3 | 3 |
| CAMK2D | 3 | 3 | 0 |
| MSSK1 | 3 | 1 | 1 |
| Haspin | 3 | 0 | 5 |
| DRAK2 | 3 | 2 | 0 |
| PAK4 | 3 | 3 | 1 |
| BMX | 3 | 2 | 1 |
| MST4 | 3 | -4 | -1 |
| CDKL1 | 2 | 3 | 1 |
| MAP2K6 | 2 | 3 | 0 |
| EPHA4 | 2 | 2 | 1 |
| MAP2K1 | 2 | 1 | 1 |
| CSNK1E | 2 | 3 | 1 |
| EPHB1 | 2 | 2 | 1 |
| BRD4 | 2 | 2 | 1 |
| PCTAIRE1 | 2 | 5 | 2 |
| CAMK1D | 2 | 0 | 0 |
| FES | 2 | -1 | 0 |
| CAMK1G | 2 | 0 | -1 |
| EPHA7 | 2 | 1 | 0 |
| CAMK2B | 2 | 2 | 1 |
| EPHA5 | 2 | 2 | 1 |
| BRPF1 | 2 | -1 | 0 |
| MST3 | 1 | -3 | -3 |

| Kinase | 2 | 6c | 7c |
|---|---|---|---|
| CAMK4 | 1 | 0 | -1 |
| STLK3 | 1 | 0 | 0 |
| MARK4 | 1 | 1 | 0 |
| PAK1 | 1 | 0 | -1 |
| FGFR1 | 1 | 0 | 0 |
| WNK1 | 1 | 2 | 0 |
| NEK1 | 1 | 0 | 0 |
| TTK | 1 | 1 | 0 |
| VRK1 | 1 | 0 | 0 |
| EphA2 | 1 | 3 | 1 |
| ATK3 | 1 | -1 | -1 |
| JNK2 | 1 | 0 | 0 |
| TLK1 | 1 | -1 | -1 |
| CASK | 0 | 1 | 0 |
| MYT1 | 0 | 0 | 0 |
| p38d | 0 | 1 | 0 |
| MAP2K7 | 0 | 0 | -1 |
| TAF1 | 0 | 1 | 1 |
| Erk2 | 0 | 0 | 0 |
| MRCKa | 0 | -1 | 0 |
| MAPKAPK2 | 0 | -1 | 0 |
| MARK3 | 0 | -1 | -1 |
| EPHB3 | 0 | 1 | -1 |
| FECH | 0 | -3 | 0 |
| FGFR2 | 0 | 0 | 0 |
| TIF1 | 0 | -2 | -1 |
| NEK7 | 0 | -1 | -1 |
| DMPK1 | 0 | 0 | 0 |
| NQO2 | -1 | -1 | -1 |
| p38a | -1 | -3 | -1 |
| OSR1 | -1 | -2 | 0 |
| NDR2 | -2 | -1 | -1 |
| NEK2 | -2 | -4 | -1 |
| **# Tm >5C** | **34** | **27** | **18** |

**Chemistry Methods**

All reagents and solvents used were purchased from commercial sources and were used without further purification. NMR spectra were obtained using a Bruker 850 MHz or INOVA 400 MHz spectrometers at room temperature; chemical shifts are expressed in parts per million (ppm, δ units) and are referenced to the residual protons in the deuterated solvent used. Coupling constants are given in units of hertz (Hz). Splitting patterns describe apparent multiplicities and are designated as s (singlet), d (doublet), t (triplet), q (quartet), m (multiplet), and br s (broad singlet), dd (doublet of doublets), ddd (double double doublet), tt (triplet of triplets). The purity of compounds submitted for biological screening was determined to be ≥95% as measured by NMR. Analytical thin layer chromatography (TLC) was performed on silica gel plates, 200 μm with an F254 indicator. Column chromatography was performed using RediSep Rf® preloaded silica gel cartridges on Isolera one Biotage automated purification systems. Samples for high-resolution mass spectrometry were analyzed with a ThermoFisher Q Exactive HF-X (ThermoFisher, Bremen, Germany) mass spectrometer coupled with a Waters Acquity H-class liquid chromatograph system. Samples were introduced via a heated electrospray source (HESI) at a flow rate of 0.3 mL/min. Electrospray source conditions were set as: spray voltage 3.0 kV, sheath gas (nitrogen) 60 arb, auxillary gas (nitrogen) 20 arb, sweep gas (nitrogen) 0 arb, nebulizer temperature 375 degrees C, capillary temperature 380 °C, RF funnel 45 V. The mass range was set to 150-2000 m/z. All measurements were recorded at a resolution setting of 120,000. Separations were conducted on a Waters Acquity UPLC BEH C18 column (2.1 x 50 mm, 1.7 $\mu$m particle size). LC conditions were set at 95 % water with 0.1% formic acid (A) ramped linearly over 5.0 mins to 100% acetonitrile with 0.1% formic acid (B) and held until 6.0 mins. At 7.0 mins the gradient was switched back to 95% (A) and allowed to re-equilibrate until 9.0 mins. Injection volume for all samples was 3 $\mu$L. Analytical LC/MS data was obtained using a Waters Acquity Ultrahigh-performance liquid chromatography (UPLC) system equipped with a photodiode array (PDA) detector using the following method: solvent A = Water + 0.2% FA, solvent B = ACN + 0.1% FA, flow rate = 1mL/min. The gradient started at 95% A for 0.05 min. Afterwards, it was ramped up to 100% B over 2 min and held for an additional minute at this concentration, before returning to the initial gradient. Compounds were purified on prep HPLC using an Agilent 1100 equipped with a Phenomenex column (Phenyl-Hexyl, 75 x 30 mm, 5 μm) using the following method: Solvent A: water + 0.05 % TFA; Solvent B: MeOH; flow rate: 70.00 mL/min. LC conditions were set at 90 % (A) ramped linearly over 8.0 mins to 100% (B) and held until 10.0 mins at 100% B. At 10.0 mins the gradient was switched back to 90% (A).

**1-(6-Chloropyrazin-2-yl)-6-nitro-1*H*-indazole (I)**

To a stirring solution of 6-nitro-1*H*-indazole (2.00 g, 1.00 Eq, 12.3 mmol) in DMF (25 mL), was added NaH (883 mg, 3 Eq, 36.8 mmol) at 0 °C. The reaction was stirred for 30 minutes at this temperature, and then 2,6-dichloropyrazine (2.74 g, 1.50 Eq, 18.4 mmol) was added. The reaction was allowed to warm to room temperature and stirred for 16 hours. The majority of DMF was removed by air drying. The resulting crude slurry was poured onto ice water and stirred for 15 minutes. Afterwards, the resulting suspension was filtered. The filter cake was washed with water several times and then left to dry *in vacuo*. The crude residue was collected and purified by silica gel chromatography using a system of hexanes and DCM starting to afford the title material as a yellowish-white solid (1.08 g, 32%). $^1$H NMR (400 MHz, $CDCl_3$) δ 9.53 (dt, *J* = 1.8, 0.8 Hz, 1H), 9.28 (d, *J* = 0.6 Hz, 1H), 8.46 (d, *J* = 0.7 Hz, 1H), 8.33 (d, *J* = 0.9 Hz, 1H), 8.16 (dd, *J* = 8.8, 2.0 Hz, 1H), 7.87 (dd, *J* = 8.8, 0.7 Hz, 1H). LCMS (ESI+) *m/z*: 276 $[M + H]^+$.

**1-(6-Chloropyrazin-2-yl)-1*H*-indazol-6-amine (II)**

To a stirring suspension of **I** (600 mg, 1 Eq, 2.18 mmol) in EtOH/water (28 mL, 3:1), was added Fe (365 mg, 3 Eq, 6.53 mmol) and $NH_4Cl$ (582 mg, 5 Eq, 10.9 mmol), and the reaction was heated to reflux and stirred for 4 hours. The reaction mixture was filtered on celite, and the celite pad was washed several times with DCM. The solvent was removed *in vacuo*. The crude residue was dissolved in DCM, and the organic layer was washed with water several times to remove excess $NH_4Cl$. The layers were separated, and the combined organic fractions were washed one more time with brine, and then dried with $Na_2SO_4$. The solvent was removed *in vacuo* to afford the title material as a yellow solid (470 mg, 88%). $^1$H NMR (400 MHz, $CDCl_3$) δ 9.26 (d, *J* = 0.6 Hz, 1H), 8.36 (d, *J* = 0.7 Hz, 1H), 8.06 (d, *J* = 0.9 Hz, 1H), 7.96 – 7.91 (m, 1H), 7.53 (dd, *J* = 8.5, 0.7 Hz, 1H), 6.72 (dd, *J* = 8.5, 2.0 Hz, 1H), 4.41 – 3.69 (br s, 2H). LCMS (ESI+) *m/z*: 246 $[M + H]^+$.

**Methyl-4-(6-(6-amino-1*H*-indazol-1-yl)pyrazin-2-yl)benzoate (III)**

A mixture of **II** (470 mg, 1 Eq, 1.91 mmol), (4-(methoxycarbonyl)phenyl)boronic acid (413 mg, 1.20 Eq, 2.30 mmol), and $Na_2CO_3$ (406 mg, 2 Eq, 3.83 mmol) were suspended in a mixture of toluene/ water/EtOH (23 mL, 4:1:1). The reaction solvent was degassed by bubbling argon gas for 5 minutes. Afterwards, $PdCl_2$(dppf) (140 mg, 0.1 Eq, 0.191 mmol) was added, and the reaction mixture was stirred under argon at 80 °C for 16 hours. The reaction mixture was poured onto water, and the aqueous layer was extracted with ethyl acetate. The combined organic extracts were washed with brine and then transferred directly to a flask. The solvent was removed *in vacuo*, and the crude product was purified using silica gel chromatography using a system of DCM/ethyl acetate to afford the title material as a yellow solid (520 mg, 78%). $^{1}$H NMR (400 MHz, DMSO-$d_6$) δ 9.25 (d, *J* = 0.6 Hz, 1H), 9.15 (d, *J* = 0.6 Hz, 1H), 8.45 (d, *J* = 8.6 Hz, 2H), 8.32 – 8.15 (m, 3H), 7.94 – 7.78 (m, 1H), 7.55 (d, *J* = 8.5 Hz, 1H), 6.71 (dd, *J* = 8.6, 1.9 Hz, 1H), 5.91 (s, 2H), 3.92 (s, 3H). LCMS (ESI+) *m/z*: 346 $[M + H]^{+}$.

**Methyl 4-(6-(5-(methylamino)-1*H*-indazol-1-yl)pyrazin-2-yl)benzoate (3a)**

A mixture of **III** (65 mg, 1 Eq, 0.19 mmol), formaldehyde (0.042 mL, 37% Wt, 3.00 Eq, 0.56 mmol), AcOH (0.011 mL, 1 Eq, 0.19 mmol) and $NaBH_3CN$ (35 mg, 3.00 Eq, 0.56 mmol) were added in DMF (1.30 mL) and stirred at room temperature overnight. Water and ethyl acetate were added. The layers were separated, and the aqueous layer was extracted with ethyl acetate. The combined organic extracts were washed with brine, filtered, and the solvent was removed *in vacuo*. The crude product was purified by silica gel chromatography using a gradient of hexanes and ethyl acetate to afford the title material as a brown solid (28 mg, 41%). $^{1}$H NMR (400 MHz, $CDCl_3$) δ 9.38 (d, *J* = 0.6 Hz, 1H), 8.87 (d, *J* = 0.6 Hz, 1H), 8.27 – 8.23 (m, 2H), 8.22 – 8.18 (m, 2H), 8.08 (d, *J* = 0.9 Hz, 1H), 8.02 (d, *J* = 2.0 Hz, 1H), 7.53 (dd, *J* = 8.7, 0.6 Hz, 1H), 6.67 (dd, *J* = 8.6, 2.1 Hz, 1H), 3.98 (s, 3H), 3.01 (s, 3H). LCMS (ESI+) *m/z*: 359 $[M + H]^{+}$.

**Methyl-4-(6-(5-(ethylamino)-1*H*-indazol-1-yl)pyrazin-2-yl)benzoate (3b)**

A mixture of methyl **III** (65 mg, 1 Eq, 0.19 mmol) and AcOH (11 mg, 1 Eq, 0.19 mmol) were added in DMF (1.30 mL). The reaction mixture was cooled to 0 °C, and then acetaldehyde (0.032 mL, 3 Eq, 0.56 mmol) and $NaBH_3CN$ (35 mg, 3.00 Eq, 0.56 mmol) were added. The reaction was quickly sealed, allowed to warm to room temperature, and then stirred overnight. Upon completion of the reaction, water and ethyl acetate were added. The layers were separated, and the aqueous layer was extracted with ethyl acetate. The combined organic extracts were washed with brine, filtered, and the solvent was removed *in vacuo*. The crude product was purified by silica gel chromatography using a gradient of hexanes and ethyl acetate to afford the title material as a brown solid (30 mg, 43%). $^{1}$H NMR (400 MHz, $CDCl_3$) δ 9.38 (d, *J* = 0.6 Hz, 1H), 8.87 (d, *J* = 0.6 Hz, 1H), 8.26 – 8.23 (m, 2H), 8.22 – 8.19 (m, 2H), 8.07 (d, *J* = 0.8 Hz, 1H), 8.03 (d, *J* = 2.0 Hz, 1H), 7.53 (dd, *J* = 8.6, 0.6 Hz, 1H), 6.65 (dd, *J* = 8.6, 2.1 Hz, 1H), 3.99 (s, 3H), 3.35 (q, *J* = 7.2 Hz, 2H), 1.39 (t, *J* = 7.1 Hz, 3H). LCMS (ESI+) *m/z*: 374 $[M + H]^{+}$.

**Methyl-4-(6-(6-(cyclopentylamino)-1H-indazol-1-yl)pyrazin-2-yl)benzoate (3c)**

Synthesized using similar procedure for **3a** starting with **III** (215 mg, 1 Eq, 0.865 mmol) and cyclopentanone (80 mg, 1.10 Eq, 0.951 mmol) to afford the title material as a yellow solid (75 mg, 15%). $^{1}$H NMR (400 MHz, $CDCl_3$) δ 9.48 (d, *J* = 0.6 Hz, 1H), 8.99 (d, *J* = 0.6 Hz, 1H), 8.27 –

8.16 (m, 4H), 8.01 (d, *J* = 2.0 Hz, 1H), 7.47 (dd, *J* = 9.0, 0.7 Hz, 1H), 6.60 – 6.56 (m, 1H), 6.52 (dd, *J* = 9.0, 1.9 Hz, 1H), 3.98 (s, 3H), 3.91 (d, *J* = 16.1 Hz, 1H), 2.11 (dq, *J* = 12.6, 6.2 Hz, 2H), 1.83 – 1.58 (m, 6H). LCMS (ESI+) *m/z*: 414 $[M + H]^+$.

**Methyl-4-(6-(6-(benzylamino)-1*H*-indazol-1-yl)pyrazin-2-yl)benzoate (3d)**

To a stirring solution of **III** (30 mg, 1 Eq, 0.087 mmol) and $NaBH_3CN$ (16 mg, 3 Eq, 0.26 mmol) in DMF (0.60 mL), were added benzaldehyde (0.018 mL, 2 Eq, 0.17 mmol) and AcOH (0.005 mL, 1 Eq, 0.087 mmol), and the resulting mixture was heated to 70 °C and stirred for 18 hours. The reaction was stopped by adding water. The aqueous layer was extracted with ethyl acetate. The combined organic extracts were washed with water, brine, and dried with $Na_2SO_4$. The solvent was removed *in vacuo*, and the crude product was purified by silica gel chromatography using a gradient of hexanes and ethyl acetate to afford the desired product as a yellow solid (20 mg, 42%). $^1$H NMR (850 MHz, DMSO-$d_6$) δ 9.24 – 9.23 (m, 1H), 9.10 – 9.10 (m, 1H), 8.31 – 8.29 (m, 2H), 8.25 (d, *J* = 0.8 Hz, 1H), 8.09 – 8.06 (m, 2H), 7.84 (d, *J* = 2.0 Hz, 1H), 7.59 (d, *J* = 8.6 Hz, 1H), 7.36 – 7.33 (m, 2H), 7.27 – 7.23 (m, 2H), 7.21 – 7.18 (m, 1H), 7.05 (t, *J* = 6.0 Hz, 1H), 6.84 (dd, *J* = 8.7, 2.0 Hz, 1H), 4.46 (d, *J* = 5.9 Hz, 2H), 3.92 (s, 3H).

**Methyl-4-(6-(6-((2-phenoxyethyl)amino)-1*H*-indazol-1-yl)pyrazin-2-yl)benzoate (3e)**

To a stirring solution of **III** (20 mg, 1.00 Eq, 0.058 mmol) and $K_2CO_3$ (16 mg, 2 Eq, 0.12 mmol) in DMF (0.50 mL), was added (2-chloroethoxy)benzene (18 mg, 2.00 Eq, 0.166 mmol), and the resulting mixture was heated to 80 °C and stirred overnight. Water was added to the reaction mixture, and the aqueous layer was extracted with ethyl acetate. The combined organic extracts were washed with water, brine, and the solvent was removed *in vacuo*. The crude product was purified by silica gel chromatography using a gradient of hexanes and ethyl acetate to afford the title material as a brown solid ( 30 mg, 40%). $^1$H NMR (850 MHz, $CDCl_3$) δ 9.38 (s, 1H), 8.87 (s, 1H), 8.24 – 8.19 (m, 4H), 8.12 (s, 1H), 8.08 (s, 1H), 7.55 (d, *J* = 8.5 Hz, 1H), 7.32 – 7.28 (m, 2H), 6.99 – 6.96 (m, 1H), 6.97 – 6.93 (m, 2H), 6.72 (dd, *J* = 8.6, 2.1 Hz, 1H), 4.56 (t, *J* = 5.8 Hz, 1H), 4.28 (t, *J* = 5.3 Hz, 2H), 3.98 (s, 3H), 3.71 (q, *J* = 5.4 Hz, 2H). LCMS (ESI+) *m/z*: 466 $[M + H]^+$.

**Methyl 4-(6-(6-((2-((*tert*-butoxycarbonyl)(methyl)amino)ethyl)amino)-1*H*-indazol-1-yl)pyrazin-2-yl)benzoate (3f)**

Synthesized using same procedure for **3d** starting with **III** (75 mg, 1 Eq, 0.22 mmol) and *tert*-butyl methyl(2-oxoethyl)carbamate (75 mg, 2 Eq, 0.43 mmol) to afford the title material as a yellow solid (47 mg, 43%). $^1$H NMR (400 MHz, DMSO-$d_6$) δ 9.26 (d, *J* = 0.7 Hz, 1H), 9.14 (d, *J* =

0.6 Hz, 1H), 8.45 – 8.37 (m, 2H), 8.25 (d, *J* = 0.7 Hz, 1H), 8.18 – 8.13 (m, 2H), 7.88 – 7.80 (m, 1H), 7.58 (d, *J* = 8.7 Hz, 1H), 6.77 (dd, *J* = 8.7, 2.0 Hz, 1H), 3.92 (s, 3H), 3.50 – 3.41 (m, 2H), 2.80 (s, 3H), 1.44 – 1.17 (m, 11H). LCMS (ESI+) *m/z*: 503 $[M + H]^+$.

**Methyl-4-(6-(6-((2-morpholinoethyl)amino)-1*H*-indazol-1-yl)pyrazin-2-yl)benzoate (3g)**

A mixture of **III** (60 mg, 1 Eq, 0.17 mmol), 4-(2-chloroethyl)morpholin-4-ium chloride (32 mg, 1.00 Eq, 0.17 mmol), KI (29 mg, 1.00 Eq, 0.17 mmol), and $K_2CO_3$ (48 mg, 2 Eq, 0.35 mmol) in ACN (2.30 mL) was heated to 170 °C under microwave conditions for 1 hour. Afterwards, water was added, and the aqueous layer was extracted with ethyl acetate. The combined organic extracts were washed with brine, and then directly transferred to a flask. The solvent was removed *in vacuo*, and the crude product was purified by silica gel chromatography using a gradient of DCM and ethyl acetate to afford the title material as a brown solid (10 mg, 12%). $^1$H NMR (400 MHz, $CDCl_3$) δ 9.38 (d, *J* = 0.6 Hz, 1H), 8.87 (d, *J* = 0.6 Hz, 1H), 8.26 – 8.23 (m, 2H), 8.21 (d, *J* = 8.8 Hz, 2H), 8.08 (d, *J* = 0.8 Hz, 1H), 8.03 (d, *J* = 2.0 Hz, 1H), 7.54 (d, *J* = 8.6 Hz, 1H), 6.72 (dd, *J* = 8.6, 2.0 Hz, 1H), 3.99 (s, 3H), 3.78 – 3.71 (m, 4H), 3.35 (t, *J* = 6.0 Hz, 2H), 2.75 (t, *J* = 6.0 Hz, 2H), 2.56 – 2.48 (m, 4H). LCMS (ESI+) *m/z*: 459 $[M + H]^+$.

**Methyl-4-(6-(6-acetamido-1*H*-indazol-1-yl)pyrazin-2-yl)benzoate (3h)**

To a stirring solution of **III** (50 mg, 1 Eq, 0.14 mmol) and DIPEA (37 mg, 0.050 mL, 2 Eq, 0.29 mmol) in THF (1.00 mL) at 0 °C, was added acetyl chloride (17 mg, 0.015 mL, 1.50 Eq, 0.22 mmol), dropwise. The reaction was stirred at 0 °C for 5 minutes, and then allowed to warm to room temperature and stirred for an additional 6 hours. Water and DCM were added. The layers were separated, and the organic layer was washed with water, brine, and then dried with $Na_2SO_4$. The solvent was removed *in vacuo*, and the crude product was purified by silica gel chromatography using a system of DCM/ethyl acetate to afford the title material as white solid (23 mg, 41%). $^1$H NMR (400 MHz, DMSO) δ 10.33 (s, 1H), 9.67 (s, 1H), 9.30 (s, 1H), 9.24 (s, 1H), 8.67 – 8.61 (m, 2H), 8.47 (d, *J* = 0.9 Hz, 1H), 8.21 – 8.10 (m, 2H), 7.85 (dd, *J* = 8.6, 0.6 Hz, 1H), 7.25 (dd, *J* = 8.6, 1.8 Hz, 1H), 3.93 (s, 3H), 2.20 (s, 3H). LCMS (ESI+) *m/z*: 387 $[M + H]^+$.

**4-(6-(6-Amino-1*H*-indazol-1-yl)pyrazin-2-yl)benzoic acid (4a)**

To a stirring solution of **III** (30 mg, 1 Eq, 0.087 mmol) in THF (0.50 mL), was added 2 M LiOH solution (0.22 mL). The reaction was heated to 50 °C and stirred overnight. Afterwards, the solvents were removed in vacuo. The residue was then dissolved in water, and the pH was adjusted to ~ 2 using 1 M HCl solution. The resulting precipitate was filtered and washed with water, and then collected. The collected residue was purified using preparative HPLC to afford the desired product as a yellow solid (15 mg, 52%). $^1$H NMR (850 MHz, DMSO) δ 9.23 (s, 1H), 9.13 (s, 1H), 8.38 (d, *J* = 8.1 Hz, 2H), 8.23 (s, 1H), 8.18 (d, *J* = 8.1 Hz, 2H), 7.87 (s, 1H), 7.55 (d, *J* = 8.4 Hz, 1H), 6.71 (dd, *J* = 8.5, 1.9 Hz, 1H), 5.91 (br s, 2H). $^{13}$C NMR (214 MHz, DMSO) δ 167.23, 150.43, 149.34, 147.83, 140.70, 138.94, 138.67, 136.26, 134.33, 133.86, 130.15, 127.00, 121.88, 117.19, 113.71, 95.61. HRMS-ESI+ (*m/z*): $[M+H]^+$ calc for $C_{18}H_{14}N_5O_2$, 332.1142; found, 332.1137.

**4-(6-(6-(Methylamino)-1*H*-indazol-1-yl)pyrazin-2-yl)benzoic acid (4b)**

Synthesized using same hydrolysis procedure for **4a** starting with **3a** (26 mg, 0.072 mmol). The dried residue was purified by reverse phase chromatography using a gradient of water/0.5%TFA and ACN to afford the TFA salt of the title material as a yellow solid (3.0 mg, 9%). $^1$H NMR (400 MHz, ) δ 13.21 (s, 1H), 9.26 (s, 1H), 9.16 (s, 1H), 8.49 – 8.36 (m, 2H), 8.25 (s, 1H), 8.14 (s, 2H), 7.82 (s, 1H), 7.57 (d, *J* = 8.7 Hz, 1H), 6.74 (dd, *J* = 8.7, 1.9 Hz, 1H), 6.53 – 6.38 (m, 1H), 2.87 (d, *J* = 3.9 Hz, 3H). $^{13}$C NMR (214 MHz, DMSO) δ 166.84, 151.09, 149.27, 147.37, 141.00, 139.59, 138.94, 136.33, 134.38, 132.01, 129.99, 126.93, 121.60, 116.85, 113.31, 92.20, 29.52. HRMS-ESI+ (*m/z*): $[M+H]^+$ calc for $C_{19}H_{16}N_5O_2$, 346.1299; found, 346.1294.

**4-(6-(6-(Ethylamino)-1*H*-indazol-1-yl)pyrazin-2-yl)benzoic acid (4c)**

To a stirring solution of **3b** (30 mg, 0.080 mmol) in THF (1.00 mL), was added 2 M LiOH solution (1.0 mL), and the reaction was stirred at room temperature overnight. The solvents were removed *in vacuo*, and the solid residue was then dissolved in water. A 1 M HCl solution was used to adjust the pH of the aqueous solution to 4-5 (or until appearance of precipitate). The suspension was filtered, and the residue was washed with water and dried to afford the title material as a yellow solid (11 mg, 38%). $^1$H NMR (400 MHz, DMSO) δ 9.25 (s, 1H), 9.14 (s, 1H), 8.42 – 8.34 (m, 2H), 8.24 (d, *J* = 0.8 Hz, 1H), 8.17 – 8.09 (m, 2H), 7.85 (s, 1H), 7.56 (d, *J* = 8.7 Hz, 1H), 6.76 (dd, *J* = 8.7, 2.0 Hz, 1H), 6.43 (t, *J* = 5.1 Hz, 1H), 3.29 – 3.20 (m, 2H), 1.29 (t, *J* = 7.1 Hz, 3H). $^{13}$C NMR (214 MHz, DMSO) δ 166.93, 150.07, 149.30, 147.45, 141.01, 139.36, 138.91, 136.32, 134.37, 132.71, 129.90, 126.86, 121.65, 116.82, 113.61, 92.28, 37.28, 14.08. HRMS-ESI+ (*m/z*): $[M+H]^+$ calc for $C_{20}H_{18}N_5O_2$, 360.1455; found, 360.1450.

**4-(6-(6-(Cyclopentylamino)-1*H*-indazol-1-yl)pyrazin-2-yl)benzoic acid (4d)**

Synthesized using same procedure for **4a** starting with **3c** (70 mg, 0.17 mmol). Purified by reverse phase chromatography using a gradient of water/0.5%TFA and ACN to afford the trifluoroacetate salt of the title material as a yellow solid ( 6 mg, 9%). $^1$H NMR (850 MHz, DMSO) δ 13.21 (br s, 1H), 9.38 – 9.26 (m, 2H), 9.17 (s, 1H), 8.52 – 8.40 (m, 2H), 8.24 – 8.03

(m, 2H), 7.50 (d, *J* = 9.0 Hz, 1H), 6.70 (dd, *J* = 9.0, 1.9 Hz, 1H), 6.30 (s, 1H), 6.13 (br s, 1H), 3.80 – 3.74 (m, 1H), 2.03 – 1.91 (m, 2H), 1.74 – 1.67 (m, 2H), 1.62 – 1.57 (m, 2H), 1.56 – 1.52 (m, 2H). $^{13}$C NMR (214 MHz, DMSO) δ 166.87, 152.81, 148.12, 148.01, 146.60, 139.39, 138.82, 134.19, 132.20, 129.88, 127.20, 121.23, 121.21, 119.85, 117.10, 53.54, 32.19, 23.92. HRMS-ESI+ (*m/z*): $[M+H]^+$ calc for $C_{23}H_{22}N_5O_2$, 400.1768; found, 400.1767.

**4-(6-(6-(Benzylamino)-1*H*-indazol-1-yl)pyrazin-2-yl)benzoic acid (4e)**

Synthesized using same procedure for **4a** starting with **3e** (20 mg, 0.046 mmol). The dried residue was purified by reverse phase chromatography using a gradient of water/0.5%TFA and ACN to afford the TFA salt of the title material as a yellow solid (3.8 mg, 20%). $^1$H NMR (400 MHz, DMSO) δ 13.18 (s, 1H), 9.24 (s, 1H), 9.10 (s, 1H), 8.32 – 8.22 (m, 3H), 8.14 – 8.04 (m, 2H), 7.84 (s, 1H), 7.59 (d, *J* = 8.6 Hz, 1H), 7.38 – 7.31 (m, 2H), 7.27 – 7.11 (m, 3H), 7.05 (t, *J* = 5.6 Hz, 1H), 6.84 (d, *J* = 8.7 Hz, 1H), 4.47 (d, *J* = 5.7 Hz, 2H). $^{13}$C NMR (214 MHz, DMSO) δ 166.85, 150.04, 149.29, 147.59, 140.82, 139.57, 139.41, 138.93, 136.52, 134.55, 131.90, 130.02, 128.27, 127.02, 127.00, 126.75, 121.79, 117.19, 113.50, 92.88, 46.29. HRMS-ESI+ (*m/z*): $[M+H]^+$ calc for $C_{25}H_{20}N_5O_2$, 422.1612; found, 422.1608.

**4-(6-(6-((2-Phenoxyethyl)amino)-1*H*-indazol-1-yl)pyrazin-2-yl)benzoic acid (4f)**

To a stirring solution of **3f** (30 mg, 1 Eq, 0.064 mmol) in THF (1 mL), was added 2 M LiOH solution (1 mL), and the reaction was heated to 50 °C and stirred for 20 hours. The solvents were evaporated. The residue was dissolved in water, and the pH was adjusted to 3 using 1 M HCl solution. The resulting suspension was filtered, and the residue was washed with water, and then collected as a yellow solid (29 mg, quantitative). $^1$H NMR (850 MHz, DMSO) δ 13.22 (s, 1H), 9.27 (s, 1H), 9.15 (s, 1H), 8.45 – 8.37 (m, 2H), 8.26 (s, 1H), 8.18 – 8.11 (m, 2H), 7.98 (s, 1H), 7.60 (d, *J* = 8.6 Hz, 1H), 7.25 (t, *J* = 7.7 Hz, 2H), 6.97 (d, *J* = 8.0 Hz, 2H), 6.90 (t, *J* = 7.3 Hz, 1H), 6.84 (d, *J* = 8.6 Hz, 1H), 6.69 (t, *J* = 5.6 Hz, 1H), 4.24 (t, *J* = 6.1 Hz, 2H), 3.71 – 3.55 (m, 2H). $^{13}$C NMR (214 MHz, DMSO) δ 166.87, 158.30, 149.87, 149.29, 147.45, 140.91, 139.56, 138.94, 136.42, 134.41, 132.03, 130.05, 129.43, 126.92, 121.80, 120.62, 117.14, 114.36, 113.67, 92.58, 65.78, 42.30. HRMS-ESI+ (*m/z*): $[M+H]^+$ calc for $C_{26}H_{22}N_5O_2$, 452.1717; found, 452.17166.

**4-(6-(6-((2-(Methylamino)ethyl)amino)-1*H*-indazol-1-yl)pyrazin-2-yl)benzoic acid (4g)**

To a stirring solution of **3g** in THF (1.00 mL) was added 2 M LiOH solution (0.40 mL), and the reaction was stirred at rt overnight . The solvents were evaporated. The residue was dissolved in water, and the pH was adjusted to 3 using 1 M HCl solution. The resulting suspension was filtered, and the residue was washed with water, and then collected. The residue was taken up in 1.20 mL of DCM/MeOH (1:4) followed by addition of TFA (0.20 mL), and the reaction was stirred at room temperature for 3 days. The solvents were removed *in vacuo*, and the crude product was purified by reverse phase chromatography using a gradient of water/0.5%TFA and ACN to afford the title material as a yellow solid (16 mg, 62%). $^{1}$H NMR (850 MHz, DMSO) δ 13.25 (s, 1H), 9.28 (s, 1H), 9.18 (s, 1H), 8.55 – 8.46 (m, 2H), 8.40 – 8.38 (m, 2H), 8.31 (s, 1H), 8.25 – 8.19 (m, 2H), 7.88 (d, *J* = 1.9 Hz, 1H), 7.67 (d, *J* = 8.6 Hz, 1H), 6.81 (dd, *J* = 8.6, 2.0 Hz, 1H), 6.49 (s, 1H), 3.51 (t, *J* = 6.1 Hz, 2H), 3.27 – 3.21 (m, 2H), 2.64 (t, *J* = 5.2 Hz, 3H). $^{13}$C NMR (214 MHz, DMSO) δ 170.10, 152.40, 152.37, 150.71, 143.77, 142.72, 142.14, 139.91, 137.81, 135.25, 133.34, 130.17, 125.23, 120.94, 116.50, 96.67, 50.10, 35.80. HRMS-ESI+ (*m/z*): $[M+H]^+$ calc for $C_{21}H_{21}N_6O_2$, 389.1721; found, 389.1721.

**4-(6-(6-((2-Morpholinoethyl)amino)-1*H*-indazol-1-yl)pyrazin-2-yl)benzoic acid (4h)**

To a stirring solution of **3h** (9.7 mg, 1 Eq, 0.021 mmol) in THF (0.80 mL) was added 2 M LiOH solution (0.80 mL), and the reaction was stirred at room temperature for 3 days . The solvents were evaporated. The residue was dissolved in water, and TFA was added until reaching a pH of 4 (or until a precipitate is formed). The resulting suspension was filtered, and the residue was washed with water, and then left to dry before being collected. The crude product was purified using preparative HPLC to afford the TFA salt of the title material as a yellow solid (3 mg, 25%). $^{1}$H NMR (400 MHz, MeOD) δ 9.25 (s, 1H), 8.93 (s, 1H), 8.34 – 8.26 (m, 2H), 8.26 – 8.18 (m, 2H), 8.12 (s, 1H), 8.01 (s, 1H), 7.60 (d, *J* = 8.6 Hz, 1H), 6.81 (d, *J* = 8.4 Hz, 1H), 3.93 – 3.82 (m, 4H), 3.68 (t, *J* = 6.1 Hz, 2H), 3.44 (t, *J* = 6.0 Hz, 2H), 3.37 – 3.29 (m, 4H). $^{13}$C NMR (214 MHz, MeOD) δ 169.22, 151.47, 150.40, 150.28, 142.41, 141.86, 139.98, 137.46, 135.82, 133.51, 131.57, 128.31, 123.20, 120.35, 114.96, 95.63, 65.03, 57.05, 53.51, 39.32. HRMS-ESI+ (*m/z*): $[M+H]^+$ calc for $C_{24}H_{25}N_6O_3$, 445.1983; found, 445.1980.

**4-(6-(6-acetamido-1*H*-indazol-1-yl)pyrazin-2-yl)benzoic acid (4i)**

Synthesized using same procedure for **4a** starting with **3i** (20 mg, 0.052 mmol) to afford the title material as a yellow solid (10 mg, 52%). $^{1}$H NMR (850 MHz, DMSO) δ 10.34 (s, 1H), 9.68 (s, 1H), 9.29 (s, 1H), 9.24 (s, 1H), 8.66 – 8.56 (m, 2H), 8.47 (s, 1H), 8.21 – 8.08 (m, 2H), 7.85 (d, *J* = 8.5 Hz, 1H), 7.26 (dd, *J* = 8.6, 1.7 Hz, 1H), 2.20 (s, 3H). $^{13}$C NMR (214 MHz, DMSO) δ 168.85, 167.02, 148.97, 147.53, 139.96, 139.09, 138.87, 138.70, 136.98, 134.50, 132.41, 129.99, 127.38, 121.71, 121.57, 116.23, 103.31, 24.43. HRMS-ESI+ (*m/z*): $[M+H]^+$ calc for $C_{20}H_{16}N_5O_3$, 374.1248; found, 374.1246.

**$N$-Isopropyl-1$H$-indazol-6-amine (IV)**

Acetone (0.055 mL, 1 Eq, 0.751 mmol), $NaBH(OAc)_3$ (318 mg, 2 Eq, 1.50 mmol) and AcOH (0.043 mL, 1 Eq, 0.751 mmol) were added to a solution of 1*H*-indazol-6-amine (100 mg, 1 Eq, 0.751 mmol) in DCM (2 mL), and the resulting mixture was stirred at room temperature for 20 hours. The reaction was terminated by adding saturated $NaHCO_3$ solution, followed by extraction with ethyl acetate. The combined organic extracts were washed with brine, dried with $Na_2SO_4$, and the solvent was removed *in vacuo*. The crude product was purified by silica gel chromatography using a gradient of DCM and ethyl acetate to afford the desired product as yellow wax (62 mg, 47%). $^1$H NMR (400 MHz, DMSO) δ 12.27 (s, 1H), 7.71 (s, 1H), 7.36 (d, *J* = 8.7 Hz, 1H), 6.50 (dd, *J* = 8.7, 1.9 Hz, 1H), 6.34 (s, 1H), 5.56 (d, *J* = 6.7 Hz, 1H), 3.60 – 3.48 (m, 1H), 1.16 (d, *J* = 6.3 Hz, 6H).

**1-(6-Iodopyrazin-2-yl)-*N*-isopropyl-1H-indazol-6-amine (V)**

(1*R*,2*R*)-*N*1,*N*2-dimethylcyclohexane-1,2-diamine (543 mg, 0.602 mL, 3.82 mmol) was added to a mixture of *N*-isopropyl-1H-indazol-6-amine (2.23 g, 1 Eq, 12.7 mmol), 2,6-diiodopyrazine (4.65 g, 1.10 Eq, 14.0 mmol), $K_3PO_4$ (6.75 g, 2.50 Eq, 31.8 mmol), and CuI (242 mg, 0.10 Eq, 1.27 mmol) in dioxane (15 mL). The mixture was purged with argon gas, sealed, heated to 110 °C and left for 24 hours. Afterwards, the reaction mixture was cooled to room temperature, diluted with ethyl acetate, and then filtered. The filtrate was concentrated *in vacuo* and the residue was purified by silica gel chromatography using a gradient of hexanes and ethyl acetate to afford the desired product as a yellow solid (2.17 g, 45%). $^1$H NMR (400 MHz, $CDCl_3$) δ 9.25 (s, 1H), 8.58 (s, 1H), 8.01 (d, *J* = 0.9 Hz, 1H), 7.76 (d, *J* = 2.0 Hz, 1H), 7.48 (d, *J* = 8.6 Hz, 1H), 6.60 (dd, *J* = 8.6, 2.0 Hz, 1H), 3.87 – 3.72 (m, 1H), 1.34 (d, *J* = 6.3 Hz, 6H). LCMS (ESI+) *m/z*: 380 $[M + H]^+$.

**4-(6-(6-(Isopropylamino)-1*H*-indazol-1-yl)pyrazin-2-yl)benzamide (5a)**

Synthesized using same procedure for **III** starting with **V** (56 mg, 1 Eq, 0.15 mmol) and (4-carbamoylphenyl)boronic acid (37 mg, 1.50 Eq, 0.22 mmol) to afford the title material as a yellow solid (32 mg, 57%). $^1$H NMR (400 MHz, DMSO) δ 9.24 (s, 1H), 9.15 (s, 1H), 8.40 – 8.32 (m, 2H), 8.22 (d, *J* = 0.8 Hz, 1H), 8.19 (br s, 1H), 8.12 – 8.05 (m, 2H), 7.85 (s, 1H), 7.60 – 7.51 (m, 2H), 6.75 (dd, *J* = 8.7, 2.0 Hz, 1H), 6.27 (d, *J* = 7.1 Hz, 1H), 3.69 (hept, *J* = 6.4 Hz, 1H), 1.31 (d, *J* = 6.3 Hz, 6H). $^{13}$C NMR (214 MHz, DMSO) δ 167.12, 149.34, 149.19, 147.58, 141.06, 138.84, 138.12, 136.27, 135.52, 134.27, 128.16, 126.57, 121.65, 116.62, 114.16, 92.29, 43.44, 22.08. HRMS-ESI+ (*m/z*): $[M+H]^+$ calc for $C_{21}H_{21}N_6O$, 373.1771; found, 373.1767.

**4-(6-(6-(Isopropylamino)-1*H*-indazol-1-yl)pyrazin-2-yl)benzenesulfonamide (5b)**

A mixture of **V** (70 mg, 1 Eq, 0.18 mmol), (4-sulfamoylphenyl)boronic acid (41 mg, 1.10 Eq, 0.20 mmol), and $Na_2CO_3$ (39 mg, 2 Eq, 0.37 mmol) were suspended in toluene/EtOH/water (4:1:1, 4.50 mL). The solvent was degassed by bubbling argon gas for 5 minutes. Afterwards, Pd $(PPh_3)_4$ (21 mg, 0.10 Eq, 0.018 mmol) was added, and the reaction mixture was stirred at 80 °C for 18 hours. Upon completion of the reaction, it was poured onto water, and the aqueous layer was extracted with ethyl acetate. The combined organic extracts were washed with brine and then dried with $Na_2SO_4$. The solvent was removed *in vacuo*, and the crude product was purified using silica gel chromatography using a gradient of hexanes and ethyl acetate to afford the title material as yellow solid (6 mg, 8%). $^1$H NMR (850 MHz, DMSO) δ 9.27 (s, 1H), 9.15 (s, 1H), 8.49 – 8.41 (m, 2H), 8.23 (s, 1H), 8.05 – 7.98 (m, 2H), 7.83 (s, 1H), 7.59 – 7.51 (m, 3H), 6.76 (dd, *J* = 8.7, 2.0 Hz, 1H), 6.29 (d, *J* = 7.1 Hz, 1H), 3.69 (hept, *J* = 6.4 Hz, 1H), 1.29 (d, *J* = 6.3 Hz, 6H). $^{13}$C NMR (214 MHz, DMSO) δ 149.34, 149.23, 147.04, 145.34, 141.03, 138.95, 138.72, 136.38, 134.67, 127.28, 126.28, 121.70, 116.65, 114.00, 92.45, 43.41, 22.12. HRMS-ESI+ (*m/z*): $[M+H]^+$ calc for $C_{20}H_{21}N_6O_2S$, 3743.1441; found, 409.1439.

**2-Fluoro-4-(6-(6-(isopropylamino)-1*H*-indazol-1-yl)pyrazin-2-yl)benzoic acid (5c)**

The ester precursor was synthesized using the same Suzuki conditions for making compound **III** starting with **V** (200 mg, 1.00 Eq, 0.527 mmol) and (3-fluoro-4-(methoxycarbonyl)phenyl)boronic acid (115 mg, 1.10 Eq, 0.580 mmol) to afford desired material as a yellow solid. The isolated product (50 mg, 0.12 mmol) was taken up in THF (1.00 mL), and then excess amount of 2 M LiOH solution was added (1 mL). The reaction was stirred at room temperature overnight. The solvents were evaporated, and the residue was triturated with DCM (x3).The residual DCM was evaporated, and the residue was then dissolved in water and the pH was adjusted 4-5 using a 1 M HCl solution. The resulting precipitate was filtered, and the residue was washed with water and dried to afford the title material as a yellow solid (30 mg, 62%). $^1$H NMR (400 MHz, DMSO) δ 13.50 (br s, 1H), 9.28 (s, 1H), 9.17 (s, 1H), 8.22 (s, 1H), 8.20 – 8.12 (m, 2H), 8.04 (t, *J* = 7.8 Hz, 1H), 7.79 (s, 1H), 7.54 (d, *J* = 8.7 Hz, 1H), 6.75 (dd, *J* = 8.7, 1.9 Hz, 1H), 6.27 (s, 1H), 3.71 (s, 1H), 1.27 (d, *J* = 6.3 Hz, 6H). $^{13}$C NMR (214 MHz, DMSO) δ 164.61, 164.60, 162.14, 160.93, 149.26 (d, $J_{C-F}$ = 11.9 Hz), 146.02, 141.65 (d, $J_{C-F}$ = 8.0 Hz), 141.04, 139.01, 136.48, 135.16, 132.59, 122.44 (d, $J_{C-F}$ = 2.9 Hz), 121.68, 120.55 (d, $J_{C-F}$ = 11.0 Hz), 116.62, 114.84 (d, $J_{C-F}$ = 24.3 Hz), 114.28, 92.23, 43.38, 22.01. HRMS-ESI+ (*m/z*): $[M+H]^+$ calc for $C_{21}H_{19}N_5O_2F$, 392.1517; found, 392.1512.

**2-Chloro-4-(6-(6-(isopropylamino)-1*H*-indazol-1-yl)pyrazin-2-yl)benzoic acid (5d)**

The ester precursor was synthesized using the same Suzuki conditions for making compound **III** staring with **V** (200 mg, 1.00 Eq, 0.527 mmol) and (3-chloro-4-(methoxycarbonyl)phenyl)boronic acid (124 mg, 1.10 Eq, 0.580 mmol) to afford the desired product as a yellow solid. The isolated product (50 mg, 0.12 mmol) was subjected to the same hydrolysis conditions as **5c** to afford the title material as a yellow solid (29 mg, 60%). $^1$H NMR (400 MHz, DMSO) δ 13.63 (br s, 1H), 9.28 (s, 1H), 9.16 (s, 1H), 8.39 (d, *J* = 1.7 Hz, 1H), 8.29 (dd, *J* = 8.2, 1.7 Hz, 1H), 8.23 (d, *J* = 0.5 Hz, 1H), 7.99 (d, *J* = 8.1 Hz, 1H), 7.79 (s, 1H), 7.54 (d, *J* = 8.7 Hz, 1H), 6.75 (dd, *J* = 8.7, 1.9 Hz, 1H), 6.26 (br s, 1H), 3.76 (hept, *J* = 6.1 Hz, 1H), 1.26 (d, *J* = 6.3 Hz, 6H). $^{13}$C NMR (214 MHz, DMSO) δ 166.28, 149.31, 149.22, 146.07, 141.05, 139.62, 139.00, 136.53, 135.07, 132.58, 132.44, 131.49, 128.35, 125.41, 121.69, 116.65, 114.45, 92.21, 43.42, 21.91. HRMS-ESI+ (*m/z*): $[M+H]^+$ calc for $C_{21}H_{19}N_5O_2Cl$, 392.1517; found, 392.1512.

**4-(6-(6-(Isopropylamino)-1*H*-indazol-1-yl)pyrazin-2-yl)-2-methoxybenzoic acid (5e)**

The ester precursor was synthesized using the same Suzuki conditions for making compound **III** staring with **V** (200 mg, 1.00 Eq, 0.527 mmol) and (3-methoxy-4-(methoxycarbonyl)phenyl)boronic acid (122 mg, 1.10 Eq, 0.580 mmol) to afford the desired product as a yellow solid. The isolated product (50 mg, 0.12 mmol) was subjected to the same hydrolysis conditions as **5c** to afford the title material as a yellow solid (17 mg, 35%). $^1$H NMR (400 MHz, DMSO) δ 9.26 (d, J = 0.5 Hz, 1H), 9.19 (s, 1H), 8.26 (s, 1H), 7.93 – 7.86 (m, 3H), 7.85 – 7.79 (m, 1H), 7.60 (d, J = 8.7 Hz, 1H), 6.80 (d, J = 8.7 Hz, 1H), 4.00 (s, 3H), 3.67 (hept, J = 6.3 Hz, 1H), 1.25 (d, J = 6.3 Hz, 6H). $^{13}$C NMR (214 MHz, DMSO) δ 170.13, 161.79, 152.40, 150.86, 143.97, 143.17, 142.02, 139.94, 137.73, 134.26, 132.77, 125.97, 124.96, 121.72, 117.22, 114.35, 59.33, 25.09. HRMS-ESI+ (*m/z*): $[M+H]^+$ calc for $C_{22}H_{22}N_5O_3$, 404.1717; found, 404.1718.

**3-Chloro-*N*-isopropyl-1*H*-indazol-6-amine (VI)**

Synthesized using same procedure for making compound **IV** starting with acetone (0.35 g, 0.44 mL, 2.00 Eq, 6.00 mmol) and 3-chloro-1*H*-indazol-6-amine (500 mg, 1.00 Eq, 3.0 mmol) to afford the title material as white solid (550 mg, 88%). $^1$H NMR (400 MHz, $CDCl_3$) δ 10.38 (s, 1H), 7.39 (d, *J* = 8.6 Hz, 1H), 6.51 (dd, *J* = 8.8, 1.9 Hz, 1H), 6.43 (d, *J* = 1.8 Hz, 1H), 3.70 – 3.59 (m, 1H), 1.26 (d, *J* = 6.3 Hz, 6H). LCMS (ESI+) *m/z*: 210 $[M + H]^+$.

**3-chloro-1-(6-iodopyrazin-2-yl)-*N*-isopropyl-1H-indazol-6-amine (VII)**

Synthesized using same procedure for making compound **V** starting with **VII** (290 mg, 1.00 Eq, 1.38 mmol) to afford the title material as a yellow solid (170 mg, 30%). $^1$H NMR (400 MHz, $CDCl_3$) δ 9.18 (d, *J* = 0.6 Hz, 1H), 8.59 (d, *J* = 0.6 Hz, 1H), 7.71 (d, *J* = 2.0 Hz, 1H), 7.42 (dd, *J* = 8.7, 0.6 Hz, 1H), 6.63 (dd, *J* = 8.8, 2.0 Hz, 1H), 4.08 (br s, 1H), 3.78 (hept, *J* = 6.3 Hz, 1H), 1.34 (d, *J* = 6.3 Hz, 6H). LCMS (ESI+) *m/z*: 413 $[M + H]^+$.

**4-(6-(3-Chloro-6-(isopropylamino)-1*H*-indazol-1-yl)pyrazin-2-yl)benzoic acid (5f)**

**VII** (0.120 g, 1.00 Eq, 0.290 mmol), (4-(methoxycarbonyl)phenyl)boronic acid (52.2 mg, 1.00 Eq, 0.290 mmol), and $Na_2CO_3$ (92.2 mg, 3.00 Eq, 0.870 mmol) were added to a mixture of toluene /EtOH/ water (3.75 mL, 4:1:1). The solvent was degassed by bubbling argon gas for 5 minutes. Then, Pd(dppf)$Cl_2$ (21.2 mg, 0.10 Eq, 0.029 mmol) was added to the reaction mixture, and the reaction flask was heated to 70 °C and stirred overnight. The reaction was cooled to room temperature and then poured onto water. The aqueous layer was extracted with ethyl acetate. The combined organic extracts were washed with brine, dried with $Na_2SO_4$, and the solvent was removed *in vacuo*. The crude product was purified by silica gel chromatography using a gradient of DCM and ethyl acetate to afford the ester precursor as a yellow solid. The isolated product (60 mg, 0.14 mmol) was subjected to the same hydrolysis conditions as **5c** to afford the title material as a yellow solid (13 mg, 22%). $^1$H NMR (850 MHz, DMSO) δ 9.13 (s, 1H), 9.10 (s, 1H), 8.27 – 8.19 (m, 2H), 8.08 – 8.04 (m, 2H), 7.82 (s, 1H), 7.45 (d, *J* = 8.7 Hz, 1H), 6.84 (dd, *J* = 8.8, 2.0 Hz, 1H), 6.58 (d, *J* = 7.0 Hz, 1H), 3.69 (hept, *J* = 6.4 Hz, 1H), 1.30 (d, *J* = 6.3 Hz, 6H). $^{13}$C NMR (214 MHz, DMSO) δ 167.33, 151.18, 150.29, 148.72, 148.24, 142.38, 138.09, 136.63, 133.41, 129.68, 126.17, 119.79, 115.24, 113.00, 91.94, 43.47, 21.97. HRMS-ESI+ (*m/z*): $[M+H]^+$ calc for $C_{21}H_{19}N_5O_2Cl$, 408.1222; found, 408.1219.

**6-Isopropoxy-1*H*-indazole (VIII)**

A mixture of 1*H*-indazol-6-ol (1.00 g, 1 Eq, 7.46 mmol), 2-iodopropane (0.97 mL, 1.30 Eq, 9.69 mmol) and $Cs_2CO_3$ (3.643 g, 1.50 Eq, 11.18 mmol) in DMF (10 mL), was stirred at room temperature for 24 hours. Water and ethyl acetate were added. The layers were separated, and the aqueous layer was extracted with ethyl acetate. The combined organic extracts were washed with brine and then dried with $Na_2SO_4$. The solvent was removed *in vacuo*, and the crude product was purified by silica gel chromatography using a gradient of hexanes and ethyl acetate to afford the desired product as a white solid (830 mg, 63%). $^1$H NMR (400 MHz, DMSO) δ 12.73 (s, 1H), 7.91 (t, *J* = 1.2 Hz, 1H), 7.59 (d, *J* = 8.7 Hz, 1H), 6.90 (s, 1H), 6.70 (dd,

*J* = 8.8, 2.1 Hz, 1H), 4.65 (hept, *J* = 6.0 Hz, 1H), 1.29 (d, *J* = 6.2 Hz, 6H). LCMS (ESI+) *m/z*: 177 $[M + H]^+$.

**1-(6-Iodopyrazin-2-yl)-6-isopropoxy-1*H*-indazole (IX)**

Synthesized using same procedure for **V** starting with **VIII** (235 mg, 1.00 Eq, 1.33 mmol) to afford the title material as white solid (158 mg, 31%). $^1$H NMR (400 MHz, $CDCl_3$) δ 9.29 (d, *J* = 0.6 Hz, 1H), 8.63 (d, *J* = 0.6 Hz, 1H), 8.15 (d, *J* = 2.2 Hz, 1H), 8.11 (d, *J* = 0.9 Hz, 1H), 7.62 (dd, *J* = 8.8, 0.6 Hz, 1H), 6.94 (dd, *J* = 8.7, 2.2 Hz, 1H), 4.73 (hept, *J* = 6.1 Hz, 1H), 1.47 (d, *J* = 6.1 Hz, 6H). LCMS (ESI+) *m/z*: 381 $[M + H]^+$.

**4-(6-(6-Isopropoxy-1*H*-indazol-1-yl)pyrazin-2-yl)benzoic acid (6a)**

The ester precursor was synthesized using similar Suzuki conditions for **5f** starting with **IX** (120 mg, 1.00 Eq, 0.316 mmol) to afford the desired material as a yellowish-white solid (75 mg, 61%). $^1$H NMR (400 MHz, DMSO) δ 9.30 (s, 1H), 9.19 (s, 1H), 8.44 (d, *J* = 0.8 Hz, 1H), 8.40 – 8.35 (m, 2H), 8.29 (d, *J* = 2.2 Hz, 1H), 8.19 – 8.13 (m, 2H), 7.80 (d, *J* = 8.8 Hz, 1H), 6.97 (dd, *J* = 8.7, 2.2 Hz, 1H), 4.76 (hept, *J* = 6.0 Hz, 1H), 3.92 (s, 3H), 1.41 (d, *J* = 6.0 Hz, 6H). LCMS (ESI+) *m/z*: 389 $[M + H]^+$. The isolated product (40 mg, 0.10 mmol) was subjected to the same hydrolysis conditions as **5c** to afford the title material as a white solid (25 mg, 65%). $^1$H NMR (400 MHz, DMSO) δ 9.25 (s, 1H), 9.15 (s, 1H), 8.43 (d, *J* = 0.8 Hz, 1H), 8.37 (d, *J* = 2.2 Hz, 1H), 8.26 – 8.20 (m, 2H), 8.11 – 8.04 (m, 2H), 7.80 (d, *J* = 8.7 Hz, 1H), 6.97 (dd, *J* = 8.8, 2.2 Hz, 1H), 4.79 (hept, *J* = 6.1 Hz, 1H), 1.44 (d, *J* = 6.0 Hz, 6H). $^{13}$C NMR (214 MHz, DMSO) δ 167.42, 158.44, 148.98, 148.44, 139.81, 138.66, 136.95, 136.68, 133.77, 129.72, 126.14, 122.46, 119.92, 115.75, 97.49, 69.92, 21.73. HRMS-ESI+ (*m/z*): $[M+H]^-$ calc for $C_{21}H_{17}N_4O_3$, 373.1301.; found, 373.1302.

**1-(6-Chloropyrazin-2-yl)-6-methoxy-1*H*-indazole (X)**

Synthesized using same procedure for **I** starting with 6-methoxy-1*H*-indazole (800 mg, 5.40 mmol) to afford the title material as a white solid (700 mg, 50%). $^1$H NMR (400 MHz, $CDCl_3$) δ 9.30 (d, *J* = 0.6 Hz, 1H), 8.40 (d, *J* = 0.6 Hz, 1H), 8.20 – 8.16 (m, 1H), 8.14 (d, *J* = 0.9 Hz, 1H), 7.64 (dd, *J* = 8.7, 0.6 Hz, 1H), 6.98 (dd, *J* = 8.7, 2.2 Hz, 1H), 3.97 (s, 3H). LCMS (ESI+) *m/z*: 261 $[M + H]^+$.

**4-(6-(6-Methoxy-1*H*-indazol-1-yl)pyrazin-2-yl)benzoic acid (6b)**

The ester precursor was synthesized using similar Suzuki conditions for **5f** starting with **X** (300 mg, 1 Eq, 1.15 mmol) and (4-(methoxycarbonyl)phenyl)boronic acid (228 mg, 1.10 Eq, 1.27

mmol) to afford the desired product as a yellow solid (134 mg, 32%). $^1H$ NMR (400 MHz, DMSO) δ 9.31 (s, 1H), 9.22 (s, 1H), 8.49 – 8.39 (m, 3H), 8.34 (br s, 1H), 8.18 (d, *J* = 8.2 Hz, 2H), 7.83 (d, *J* = 8.8 Hz, 1H), 7.03 (dd, *J* = 8.9, 2.2 Hz, 1H), 3.97 (s, 3H), 3.92 (s, 3H). LCMS (ESI+) *m/z*: 361 $[M + H]^+$. The isolated product (50 mg, 0.14 mmol) was subjected to the same hydrolysis conditions as **5c** to afford the title material as a white solid (25 mg, 52%). $^1H$ NMR (400 MHz, DMSO) δ 13.20 (br s, 1H), 9.29 (d, *J* = 0.4 Hz, 1H), 9.20 (d, *J* = 0.6 Hz, 1H), 8.45 (d, *J* = 0.8 Hz, 1H), 8.41 – 8.36 (m, 2H), 8.34 (d, *J* = 2.2 Hz, 1H), 8.20 – 8.07 (m, 2H), 7.82 (d, *J* = 8.7 Hz, 1H), 7.02 (dd, *J* = 8.8, 2.2 Hz, 1H), 3.96 (s, 3H). $^{13}C$ NMR (214 MHz, DMSO) δ 166.85, 160.34, 148.93, 147.61, 139.78, 139.40, 138.84, 137.23, 134.44, 132.21, 130.09, 126.99, 122.40, 120.12, 114.66, 96.17, 55.32. HRMS-ESI+ (*m/z*): $[M+H]^+$ calc for $C_{19}H_{15}N_4O_3$, 347.1139.; found, 347.1134.

**Methyl- 4-(6-chloropyrazin-2-yl)benzoate (XI)**

2-bromo-6-chloropyrazine (2.15 g, 1 Eq, 11.1 mmol), (4-(methoxycarbonyl)phenyl)boronic acid (2.00 g, 1 Eq, 11.1 mmol) and $Na_2CO_3$ (2.36 g, 2 Eq, 22.2 mmol) were suspended in a mixture of toluene/EtOH/water (60 mL, 4:1:1). The solvent was degassed by bubbling argon gas for 10 minutes, and then $Pd(PPh_3)_4$ (1.03 g, 0.08 Eq, 0.889 mmol) was added to the reaction mixture and it was heated to 70 °C and stirred for 6 hours. Afterwards, the reaction mixture was poured into water, and the aqueous layer was extracted with ethyl acetate. The combined organic extracts were dried over anhydrous $Na_2SO_4$, filtered, and then concentrated *in vacuo.* The crude product was purified by silica gel chromatography using a gradient of hexanes and ethyl acetate to afford the title material as a white solid (1.36 g, 49%). $^1H$ NMR (400 MHz, $CDCl_3$) δ 8.98 (s, 1H), 8.58 (s, 1H), 8.21 – 8.15 (m, 2H), 8.14 – 8.07 (m, 2H), 3.96 (s, 3H). LCMS (ESI+) *m/z*: 249 $[M + H]^+$.

**4-(6-(6-Isopropoxy-1*H*-indol-1-yl)pyrazin-2-yl)benzoic acid (6c)**

To a solution of 6-isoproxyindole (190 mg, 1.00 Eq, 1.08 mmol) in DMF (2.00 mL), was added NaH (78.1 mg, 3.00 Eq, 3.25 mmol) at 0 °C. The reaction was stirred for 30 minutes, and then **XI** (297 mg, 1.10 Eq, 1.19 mmol) in DMF (3.50 mL) was added portion-wise. The reaction was allowed to warm to room temperature and then stirred for 18 hours. It was noticed that the ester was hydrolyzed in the reaction, so 1 M HCl solution was added to the reaction mixture to ensure protonation of the carboxylic acid. Ethyl acetate was added, the layers were separated, and the aqueous layer was extracted with ethyl acetate. The combined organic extracts were washed with brine, dried with $Na_2SO_4$, and then filtered. The solvent was removed *in vacuo*. The crude product was purified by silica gel chromatography using a system of DCM/MeOH followed by reverse phase chromatography (water/TFA 0.5% and ACN) to afford the trifluoroacetate salt of the title material as a yellowish white solid (25 mg, 10%). $^{1}$H NMR (850 MHz, DMSO) δ 13.23 (br s, 1H), 9.18 (s, 1H), 9.17 (s, 1H), 8.38 – 8.35 (m, 2H), 8.25 (d, *J* = 2.3 Hz, 1H), 8.15 (d, *J* = 3.6 Hz, 1H), 8.15 – 8.11 (m, 2H), 7.55 (d, *J* = 8.5 Hz, 1H), 6.86 (dd, *J* = 8.5, 2.3 Hz, 1H), 6.79 (dd, *J* = 3.5, 0.6 Hz, 1H), 4.65 (hept, *J* = 6.1 Hz, 1H), 1.34 (d, *J* = 6.0 Hz, 6H). $^{13}$C NMR (214 MHz, DMSO) δ 166.84, 154.90, 148.26, 147.90, 139.61, 136.89, 135.61, 135.25, 132.11, 129.90, 126.99, 124.75, 124.02, 121.51, 113.43, 107.00, 100.49, 69.85, 21.97. HRMS-ESI+ (*m/z*): $[M+H]^+$ calc for $C_{22}H_{20}N_3O_3$, 374.1499.; found, 374.1499.

**4-(6-(6-(Isopropylamino)-1*H*-indol-1-yl)pyrazin-2-yl)benzoic acid (6d)**

The ester precursor was synthesized using similar procedure for **V** starting with **XI** (320 mg, 1 Eq, 1.29 mmol) and *N*-isopropyl-1*H*-indol-6-amine (247 mg, 1.10 Eq, 1.42 mmol) to afford the desired product as a yellow solid. The isolated product (60 mg, 1 Eq, 0.16 mmol) was subjected to the same hydrolysis conditions as **5c** to afford the title material a white solid (10 mg, 17%). $^{1}$H NMR (850 MHz, MeOD) δ 9.06 (s, 1H), 9.01 (s, 1H), 8.48 (s, 1H), 8.34 – 8.32 (m, 2H), 8.25 – 8.22 (m, 2H), 8.08 (d, *J* = 3.6 Hz, 1H), 7.72 (d, *J* = 8.3 Hz, 1H), 7.10 (dd, *J* = 8.3, 2.0 Hz, 1H), 6.86 (d, *J* = 3.3 Hz, 1H), 3.79 (hept, *J* = 6.4 Hz, 1H), 1.37 (d, *J* = 6.4 Hz, 6H). $^{13}$C NMR (214 MHz, MeOD) δ 169.23, 150.93, 150.13, 141.34, 138.34, 136.97, 135.96, 133.68, 131.49, 129.73, 128.40, 127.15, 123.45, 116.60, 108.41, 20.59.

**Methyl-4-(6-((2-fluoro-5-nitrophenyl)amino)pyrazin-2-yl)benzoate (XIIa)**

**XI** (145 mg, 1 Eq, 0.583 mmol), 2-fluoro-5-nitroaniline (137 mg, 1.50 Eq, 0.875 mmol), $Cs_2CO_3$ (380 mg, 2 Eq, 1.17 mmol), BINAP (36.3 mg, 0.10 Eq, 0.058 mmol), and $Pd_2(dba)_3$ (26.7 mg, 0.05 Eq, 0.029 mmol) were mixed in dioxane (3.50 mL). The solvent was purged with argon gas, and then the flask was sealed. The reaction mixture was heated to 90 °C and stirred for 16 hours. Upon completion of the reaction, it was allowed to cool to room temperature, diluted with DCM, and then filtered over a celite pad. The residue on the celite pad was washed several times with DCM. The filtrate was collected and concentrated *in vacuo*. The crude compound was purified using silica gel chromatography using a gradient of hexanes and ethyl acetate to afford the desired product as a yellow solid (95 mg, 44%). $^{1}$H NMR (400 MHz, $cdcl_3$) δ 9.89 (dd, *J* = 7.2, 2.8 Hz, 1H), 8.71 (d, *J* = 0.6 Hz, 1H), 8.27 (d, *J* = 0.6 Hz, 1H), 8.25 – 8.20 (m, 4H), 7.94 (ddd, *J* = 8.9, 4.3, 2.8 Hz, 1H), 7.29 (dd, *J* = 10.3, 9.0 Hz, 1H), 7.02 (d, *J* = 4.0 Hz, 1H), 3.97 (s, 3H). LCMS (ESI+) *m/z*: 369 $[M + H]^+$.

**Methyl-4-(6-((2-methoxy-5-nitrophenyl)amino)pyrazin-2-yl)benzoate (XIIb)**
Synthesized using same procedure as **XIIa** starting with **XI** (300 mg, 1 Eq, 1.21 mmol) and 2-methoxy-5-nitroaniline (304 mg, 1.50 Eq, 1.81 mmol) to afford the title material as a yellow solid (250 mg, 55%). $^{1}$H NMR (850 MHz, DMSO) δ 9.81 – 9.71 (m, 1H), 9.31 (d, *J* = 2.7 Hz, 1H), 8.79 – 8.73 (m, 1H), 8.65 (s, 1H), 8.41 – 8.30 (m, 2H), 8.12 – 8.04 (m, 2H), 7.99 – 7.92 (m, 1H), 7.28 (dd, *J* = 9.0, 1.5 Hz, 1H), 4.07 (s, 3H), 3.90 (s, 3H). LCMS (ESI+) *m/z*: 381 $[M + H]^{+}$.

**Methyl 4-(6-((5-amino-2-fluorophenyl)amino)pyrazin-2-yl)benzoate (XIIIa)**
To a stirring suspension of **XIIa** (90 mg, 1 Eq, 0.24 mmol) in 3:1 EtOH/Water (3.50 mL), were added Fe (68 mg, 5 Eq, 1.2 mmol) and $NH_4Cl$ (39 mg, 3 Eq, 0.73 mmol), and the reaction was heated to reflux and stirred for 3 hours. The reaction was stopped and filtered through celite. The filtering agent was washed several times with ethyl acetate. The filtrate was concentrated *in vacuo*. The crude product was dissolved in Ethyl acetate. Water was added, and the organic phase was washed with water and brine and then dried with $Na_2SO_4$. The solvent was removed *in vacuo* to afford the desired amine as a green solid which was carried to the next step without further purification.

**Methyl-4-(6-((5-amino-2-methoxyphenyl)amino)pyrazin-2-yl)benzoate (XIIIb)**
Synthesized using similar procedure to **XIIIa** starting with **XIIa** (200 mg, 0.526 mmol) to afford the desired amine as a yellow solid which was carried to the next step without further purification.

**4-(6-((2-Fluoro-5-(isopropylamino)phenyl)amino)pyrazin-2-yl)benzoic acid (7a)**
Acetone (20 mg, 0.025 mL, 3.00 Eq, 0.34 mmol),**XIIIa** (38 mg, 1 Eq, 0.11 mmol) and AcOH (0.006 mL, 1 Eq, 0.11 mmol) were mixed in DCM (2.50 mL), and the resulting mixture was stirred at room temperature for 10 minutes followed by the addition of $NaBH(OAc)_3$ (48 mg, 2.00 Eq, 0.22 mmol). The reaction was stirred at room temperature overnight. An additional 1 mL of acetone was added, and the reaction was stirred for an additional 4 hours. The reaction was terminated by the use of a saturated $NaHCO_3$ solution, followed by extraction with ethyl acetate. The solvent was removed *in vacuo*. The crude product was purified by a silica gel chromatography using a gradient of hexanes and ethyl acetate to afford the desired product as brown solid. The isolated product (10 mg, 0.026 mmol) was subjected to the same hydrolysis conditions as **5c** to afford the title material as a yellow solid (9 mg, 93%). $^{1}$H NMR (850 MHz, MeOD) δ 8.49 (s, 1H), 8.21 (s, 1H), 8.18 (d, *J* = 8.4 Hz, 2H), 8.12 (d, *J* = 8.5 Hz, 2H), 7.78 (dd, *J* = 7.0, 2.8 Hz, 1H), 6.95 (dd, *J* = 10.9, 8.7 Hz, 1H), 6.43 – 6.34 (m, 1H), 3.62 (hept, *J* = 6.3 Hz,

1H), 1.22 (d, *J* = 6.3 Hz, 6H). $^{13}$C NMR (214 MHz, MeOD) δ 169.71, 153.72, 149.90, 149.03, 147.93, 145.71, 142.26, 134.50, 133.27, 131.54, 131.13, 129.47 (d, $J_{C\text{-}F}$ = 11.8 Hz), 127.88, 116.30, 110.66 (d, $J_{C\text{-}F}$ = 7.0 Hz), 108.52, 46.69, 22.88. HRMS-ESI+ (*m/z*): $[M+H]^+$ calc for $C_{20}H_{20}N_4O_2F$, 367.1565.; found, 367.1563.

**4-(6-((5-(Isopropylamino)-2-methoxyphenyl)amino)pyrazin-2-yl)benzoic acid (7b)**

The ester precursor was synthesized using similar procedure to **7a** starting with **XIIIb** (180 mg, 1 Eq, 0.526 mmol) to afford the desired product as brown solid. The isolated product (93 mg, 0.24 mmol) was subjected to the same hydrolysis conditions as **5c.** After ester hydrolysis, the product was purified by reverse phase HPLC chromatography to afford the trifluoroacetate salt of the title material as a yellow solid (17 mg, 19%).$^{1}$H NMR (850 MHz, DMSO) δ 13.16 (s, 1H), 10.44 (s, 2H), 9.12 (s, 1H), 8.73 – 8.57 (m, 3H), 8.27 – 8.21 (m, 2H), 8.10 – 8.01 (m, 2H), 7.21 (d, *J* = 8.5 Hz, 1H), 7.15 – 6.96 (m, 1H), 3.95 (s, 3H), 3.67 (hept, *J* = 6.5 Hz, 1H), 1.30 (d, *J* = 6.4 Hz, 6H). $^{13}$C NMR (214 MHz, DMSO) δ 166.94, 151.13, 146.49, 140.37, 135.65, 131.61, 131.48, 129.98, 129.71, 126.87, 111.53, 56.26, 18.87. HRMS-ESI+ (*m/z*): $[M+H]^+$ calc for $C_{21}H_{23}N_4O_3$, 367.1765.; found, 379.1763.

**4-Isopropoxy-1-methoxy-2-nitrobenzene (XIV)**

A mixture of 4-methoxy-3-nitrophenol (500 mg, 1 Eq, 2.96 mmol), 2-iodopropane (653 mg, 0.384 mL, 1.30 Eq, 3.84 mmol) and $Cs_2CO_3$ (1.44 g, 1.50 Eq, 4.43 mmol) in DMF (5.00 mL) was stirred at room temperature for 24 hours, and at 60 °C for 4 hours . Water was added and the aqueous layer was extracted with ethyl acetate. The combined organic extracts were washed with brine and then dried with $Na_2SO_4$. The solvent was removed *in vacuo*. The crude product was purified by silica gel chromatography using a gradient of hexanes and ethyl acetate to afford the desired product as a yellow liquid (454 mg, 73%). $^{1}$H NMR (400 MHz, $CDCl_3$) δ 7.39 (d, *J* = 3.0 Hz, 1H), 7.09 (dd, *J* = 9.1, 3.0 Hz, 1H), 7.01 (d, *J* = 9.1 Hz, 1H), 4.48 (hept, *J* = 6.0 Hz, 1H), 3.91 (s, 3H), 1.33 (d, *J* = 6.1 Hz, 6H). LCMS (ESI+) *m/z*: 212 $[M + H]^+$.

**5-Isopropoxy-2-methoxyaniline (XVI)**

Synthesized using same procedure for **XIIIa** starting with **XIV** (400 mg, 1.89 mmol) to afford the title material in quantitative yield and taken to the next step without purification.

**4-(6-((5-Isopropoxy-2-methoxyphenyl)amino)pyrazin-2-yl)benzoic acid (7c)**

The ester precursor was synthesized using same procedure for **XIIa** starting with **XI** (250 mg, 1 Eq, 1.01 mmol) and **XVI** (219 mg, 1.20 Eq, 1.21 mmol) to afford the desired product as a yellow solid. The isolated product (100 mg, 1 Eq, 0.254 mmol) was subjected to the same hydrolysis conditions as **5c** to afford the title material as a yellow solid (90 mg, 93%). $^1$H NMR (850 MHz, DMSO) δ 13.12 (s, 1H), 8.83 (s, 1H), 8.60 (s, 1H), 8.49 (s, 1H), 8.21 – 8.18 (m, 2H), 8.15 (d, *J* = 3.0 Hz, 1H), 8.06 – 7.99 (m, 2H), 6.96 (d, *J* = 8.7 Hz, 1H), 6.56 (dd, *J* = 8.8, 3.0 Hz, 1H), 4.50 (hept, *J* = 6.0 Hz, 1H), 3.84 (s, 3H), 1.27 (d, *J* = 6.1 Hz, 6H). $^{13}$C NMR (214 MHz, DMSO) δ 166.92, 151.47, 151.19, 146.39, 143.44, 140.68, 135.32, 131.39, 130.79, 129.63, 126.48, 111.86, 109.85, 108.26, 70.32, 56.23, 22.07. HRMS-ESI+ (*m/z*): $[M+H]^+$ calc for $C_{21}H_{22}N_3O_4$, 380.1605.; found, 380.1604.

### Biology Methods

**NanoBRET Assay.** Assays were run with a modified version of the previously published protocols.[5, 8] HEK293 cells were cultured at 37 °C, 5% $CO_2$ in Dulbecco's modified Eagle medium (DMEM; Gibco) supplemented with 10% fetal bovine serum (VWR/Avantor). A transfection complex of DNA at 10 μg/mL was created, consisting of 9 μg/mL of carrier DNA (Promega) and 1 μg/mL of CSNK2A2-NLuc fusion DNA in Opti-MEM without serum (Gibco). FuGENE HD (Promega) was added at 30 μl/mL to form a lipid:DNA complex. The solution was then mixed and incubated at room temperature for 20 min. The transition complex was mixed with a 20x volume of HEK293 cells at 20,000 cells per mL in DMEM/FBS and 100 μL per well was added to a 96-well plate that was incubated overnight at 37°C, 5% $CO_2$. The following day, the media was removed via aspiration and replaced with 85 μL of Opti-MEM without phenol red. A total of 5 μL per well of 20x-NanoBRET Tracer K10 (Promega) at 5 μM in Tracer Dilution Buffer (Promega N291B) was added to all wells, except the "no tracer" control wells. Test compounds (10 mM in DMSO) were diluted 100x in Opti-MEM media to prepare stock solutions and evaluated at eleven concentrations. A total of 10 μL per well of the 10-fold test compound stock solutions (final assay concentration of 0.1% DMSO) were added. For "no compound" and "no tracer" control wells, DMSO in OptiMEM was added for a final concentration of 1.1% across all wells. 96-well plates containing cells with NanoBRET Tracer K10 and test compounds (100 μL total volume per well) were equilibrated (37°C / 5% $CO_2$) for 2 h. The plates were cooled to room temperature for 15 min. NanoBRET NanoGlo substrate (Promega) at a ratio of 1:166 to Opti-MEM media in combination with extracellular NLuc Inhibitor (Promega) diluted 1:500 (10 μL of 30 mM stock per 5 mL Opti-MEM plus substrate) were combined to create a 3X stock solution. A total of 50 μL of the 3X substrate/extracellular NL inhibitor were added to each well. The plates were read within 30 min on a GloMax Discover luminometer (Promega) equipped with 450 nm BP filter (donor) and 600 nm LP filter (acceptor) using 0.3 s integration time. Raw milliBRET (mBRET) values were obtained by dividing the acceptor emission values (600 nm) by the donor emission values (450 nm) and multiplying by 1000. Averaged control values were used to represent complete inhibition (no tracer control: Opti-MEM + DMSO only) and no inhibition (tracer only control: no compound, Opti-MEM + DMSO + Tracer K10 only) and were plotted alongside the raw mBRET values. The data was first normalized and then fit using Sigmoidal, 4PL binding curve in Prism Software to determine $IC_{50}$ values.

**MHV Assay.** DBT cells were cultured at 37°C in Dulbecco's modified Eagle medium (DMEM; Sigma) supplemented with 10% fetal bovine serum (Gibco) and penicillin and streptomycin (Sigma). DBT cells were plated in 96 well plates to be 80% confluent at the start of the assay. Test compounds were diluted to 15 µM in DMEM. Serial 4-fold dilutions were made in DMEM, providing a concentration range of 15 µM to 0.22 µM. Media was aspirated from the DBT cells and 100 µL of the diluted test compounds were added to the cells for 1 h at 37°C. After 1 h, MHV-nLuc[5] was added at an MOI of 0.1 in 50 µL DMEM so that the final concentration of the first dilution of compound was 10 µM (T=0). After 10 h, the media was aspirated, and the cells were washed with PBS and lysed with passive lysis buffer (Promega) for 20 min at room temperature. Relative light units (RLUs) were measured using a luminometer (Promega; GloMax). Triplicate data was analyzed in Prism Graphpad to generate $IC_{50}$ values.

**Protein Expression and purification.** Expression and purification were performed as described previously[1-3]. Briefly, transformed BL21(DE3) cells were grown in Terrific Broth medium containing 50 mg/mL kanamycin. Protein expression was induced at an OD600 of 2 by using 0.5 mM isopropyl-thio-galactopyranoside (IPTG) at 18 C for 12 hours. Cells expressing His6-tagged CSNK2A1 were lysed in lysis buffer containing 50 mM HEPES pH 7.5, 500 mM NaCl, 25 mM imidazole, 5% glycerol, and 0.5 mM Tris(2 carboxyethyl)phosphine (TCEP) by sonication. After centrifugation, the supernatant was loaded onto a Nickel-Sepharose column equilibrated with 30 mL lysis buffer. The column was washed with 60 mL lysis buffer. Proteins were eluted by an imidazole step gradient (50, 100, 200, 300 mM). Fractions containing protein were pooled together and dialyzed overnight using 1L of final buffer (25 mM HEPES pH 7.5, 500 mM NaCl, 0.5 mM TCEP) at 4 C. Additionally, TEV protease was added (protein:TEV 1:20 molar ratio) to remove the tag. The next day the protein solution was loaded onto Nickel-Sepharose column beads again to remove the TEV protease and cleaved Tag. The combined flow through fraction and the wash fraction (25 mM imidazole) containing the protein were concentrated to approximately 4-5 mL and loaded onto Superdex 75 16/60 Hi-Load gel filtration column equilibrated with final buffer. The protein was concentrated to approximately 9 mg/mL.

**Crystallization.** CSNK2A1 was crystallized using the sitting-drop vapor diffusion method by mixing protein (9 mg/mL) and well solutions in 2:1, 1:1, and 1:2 ratios. The reservoir solution contained 0.2 M ammonia sulfate, 0.1 M bis-tris pH 5.5 and between 23-26% (v/v) PEG 3350. Complex structures were achieved by soaking the apo crystals for at least 24h with the desired inhibitor dissolved in reservoir solution. Final concentration of the inhibitor was 0.5 mM.

**Data collection, structure solution and refinement.** Diffraction data were collected at beamline X06SA (Villigen, CH) at a wavelength of 1.0 Å at 100 K. The reservoir solution supplemented with 20% ethylene glycol was used as cryoprotectant. Data were processed using XDS[4] and scaled with aimless[5]. The PDB structure with the accession code 6Z83[1] was used as an initial search MR model using the program MOLREP[6]. The final model was built manually using Coot[7] and refined with REFMAC5[8]. Data collection and refinement statistics:

| **Data collection** | **CSNK2A1-7c** | **CSNK2A1-6c** |
|---|---|---|
| Beamline | X06SA/PXI SLS | X06SA/PXI SLS |
| Wavelength (Å) | 1.00000 | 1.00000 |
| Space group | $P4_32_12$ | $P4_32_12$ |
| Cell dimensions | | |
| *a, b, c (Å)* | 127.97, 127.97, 125.13 | 127.84, 127.84, 125.34 |
| α, β, γ (°) | 90, 90, 90 | 90, 90, 90 |
| Resolution (Å)* | 45.24-2.60 (2.72-2.60) | 45.20-2.60 (2.72-2.60) |
| unique observations* | 32597 (3927) | 32597 (3924) |
| *Rpim** | 0.06 (0.77) | 0.03 (0.54) |
| Completeness (%)* | 99.9 (99.9) | 99.9 (99.9) |
| Multiplicity* | 13.2 (13.6) | 13.3 (13.7) |
| mean I/σI* | 10.9 (1.8) | 15.9 (1.9) |
| CC1/2* | 0.99 (0.77) | 0.99 (0.76) |
| **Refinement** | | |
| *Rwork / Rfree* | 0.2105 / 0.2509 | 0.2053 / 0.2457 |
| No. of atoms | 5449 | 5470 |
| overall B-factors ($Å^2$) | 69.238 | 73.395 |
| Rms deviations | | |
| Bond lengths (Å) | 0.0076 | 0.0069 |
| Bond angles (°) | 1.4537 | 1.4665 |
| Ramachandran outlier (%) | 0.0 | 0.0 |
| **Protein Data Bank entry** | 8QWZ | 8QWY |

*Values for the highest resolution shell are shown in parentheses.

**Thermal stability kinase selectivity.** The assay was performed as previously described.[9,10] Briefly, recombinant protein kinase domains at a concentration of 2 µM were mixed with 10 µM compound in a buffer containing 20 mM HEPES, pH 7.5, and 500 mM NaCl. SYPRO Orange (5000×, Invitrogen) was added as a fluorescence probe (1 µl per mL). Subsequently, temperature-dependent protein unfolding profiles were measured using the QuantStudio™ 5 realtime PCR machine (Thermo Fisher). Excitation and emission filters were set to 465 nm and 590 nm, respectively. The temperature was raised with a step rate of 3°C per minute. Data points were analysed with the internal software (Thermal Shift Software™ Version 1.4, Thermo Fisher) using the Boltzmann equation to determine the inflection point of the transition curve.

### References


1. Wells, C. I.; Drewry, D. H.; Pickett, J. E.; Tjaden, A.; Krämer, A.; Müller, S.; Gyenis, L.; Menyhart, D.; Litchfield, D. W.; Knapp, S.; Axtman, A. D., Development of a potent and selective chemical probe for the pleiotropic kinase CK2. *Cell Chemical Biology* **2021**.
2. Davis-Gilbert, Z. W.; Krämer, A.; Dunford, J. E.; Howell, S.; Senbabaoglu, F.; Wells, C. I.; Bashore, F. M.; Havener, T. M.; Smith, J. L.; Hossain, M. A.; Oppermann, U.; Drewry, D. H.; Axtman, A. D., Discovery of a Potent and Selective Naphthyridine-Based Chemical Probe for Casein Kinase 2. *ACS Medicinal Chemistry Letters* **2023**.
3. Krämer, A.; Kurz, C. G.; Berger, B.-T.; Celik, I. E.; Tjaden, A.; Greco, F. A.; Knapp, S.; Hanke, T., Optimization of pyrazolo[1,5-a]pyrimidines lead to the identification of a highly selective casein kinase 2 inhibitor. *European Journal of Medicinal Chemistry* **2020,** *208*, 112770.
4. Kabsch, W., XDS. *Acta Crystallographica Section D: Biological Crystallography* **2010,** *66* (Pt 2), 125-132.
5. Evans, P. R.; Murshudov, G. N., How good are my data and what is the resolution? *Acta Crystallographica Section D: Biological Crystallography* **2013,** *69* (Pt 7), 1204-1214.
6. Lebedev, A. A.; Vagin, A. A.; Murshudov, G. N., Model preparation in MOLREP and examples of model improvement using X-ray data. *Acta Crystallographica Section D: Biological Crystallography* **2008,** *64* (Pt 1), 33-39.
7. Emsley, P.; Cowtan, K., Coot: model-building tools for molecular graphics. *Acta Crystallogr D Biol Crystallogr* **2004,** *60* (Pt 12 Pt 1), 2126-32.
8. Vagin, A. A.; Steiner, R. A.; Lebedev, A. A.; Potterton, L.; McNicholas, S.; Long, F.; Murshudov, G. N., REFMAC5 dictionary: organization of prior chemical knowledge and guidelines for its use. *Acta Crystallogr D Biol Crystallogr* **2004,** *60* (Pt 12 Pt 1), 2184-95.
9. Krämer, A.; Kurz, C. G.; Berger, B.-T.; Celik, I. E.; Tjaden, A.; Greco, F. A.; Knapp, S.; Hanke, T. Optimization of pyrazolo[1,5-a]pyrimidines lead to the identification of a highly selective casein kinase 2 inhibitor. *European Journal of Medicinal Chemistry* **2020,** 208, 112770.
10. Fedorov, O.; Niesen, F. H.; Knapp, S. Kinase inhibitor selectivity profiling using differential scanning fluorimetry. *Methods Mol Biol* **2012,** 795, 109-18.
